# Nicotinamide riboside augments the human skeletal muscle NAD^+^ metabolome and induces transcriptomic and anti-inflammatory signatures in aged subjects: a placebo-controlled, randomized trial

**DOI:** 10.1101/680462

**Authors:** Yasir S Elhassan, Katarina Kluckova, Rachel S Fletcher, Mark Schmidt, Antje Garten, Craig L Doig, David M Cartwright, Lucy Oakey, Claire V Burley, Ned Jenkinson, Martin Wilson, Samuel J E Lucas, Ildem Akerman, Alex Seabright, Yu-Chiang Lai, Daniel A Tennant, Peter Nightingale, Gareth A Wallis, Konstantinos N Manolopoulos, Charles Brenner, Andrew Philp, Gareth G Lavery

**Author notes:** Correspondence to: Gareth G Lavery Office 231, IBR Tower, Institute of Metabolism and Systems Research, University of Birmingham, Edgbaston, B15 2TT, UK.

## Abstract

NAD^+^ is modulated by conditions of metabolic stress and has been reported to decline with aging, but human data are sparse. Nicotinamide riboside (NR) supplementation ameliorates metabolic dysfunction in rodents. We aimed to establish whether oral NR supplementation in aged participants can increase the skeletal muscle NAD^+^ metabolome, and questioned if tissue NAD^+^ levels are depressed with aging. We supplemented 12 aged men with NR 1g per day for 21-days in a placebo-controlled, randomized, double-blind, crossover trial. Targeted metabolomics showed that NR elevated the muscle NAD^+^ metabolome, evident by increased nicotinic acid adenine dinucleotide and nicotinamide clearance products. Muscle RNA sequencing revealed NR-mediated downregulation of energy metabolism and mitochondria pathways. NR also depressed levels of circulating inflammatory cytokines. In an additional study, ^31^P magnetic resonance spectroscopy-based NAD^+^ measurement in muscle and brain showed no difference between young and aged individuals. Our data establish that oral NR is available to aged human muscle and identify anti-inflammatory effects of NR, while suggesting that NAD^+^ decline is not associated with chronological aging *per se* in human muscle or brain.

## INTRODUCTION

Aging is characterized by a decline in metabolic and physiological functions of all organs within the body. A hallmark feature of aging is the progressive loss of skeletal muscle mass and function that can progress to sarcopenia, which is associated with significant morbidity, mortality, and substantial healthcare costs (Kim and Choi, 2013; Sousa *et al*., 2016). Exercise is considered a front line modality to combat age-related muscle decline (Costford *et al*., 2010). However, nutritional strategies may also offer an effective countermeasure to age-associated morbidities and promote healthy muscle aging (Bogan and Brenner, 2008). Nicotinamide adenine dinucleotide (NAD^+^) homeostasis is critical to cell and organismal function. As well as its classical role in redox metabolism, NAD^+^ is a substrate for enzymes such as sirtuins, poly-ADPribose polymerases (PARPs), and cyclic ADPribose synthetases that regulate key cellular processes of energy metabolism, DNA damage repair, and calcium signalling (Yoshino, Baur and Imai, 2017). Improving NAD^+^ availability via the supplementation of the NAD^+^ precursor nicotinamide riboside (NR) (Bieganowski and Brenner, 2004; Trammell *et al*., 2016) has emerged as a potential strategy to augment tissue-specific NAD^+^ homeostasis, and improve physiological function (Elhassan, Philp and Lavery, 2017). A range of physiological stresses associated with depletion of NAD^+^ and/or NADPH have been ameliorated with NR supplementation in mice including prevention of noise-induced hearing loss (Brown *et al*., 2014), resistance to weight gain (Cantó *et al*., 2012), reduction of blood glucose, hepatic steatosis, and neuropathy on high fat diet (Trammell et al. 2016), improvement of cardiac function in genetic cardiomyopathy (Diguet *et al*., 2018), and prevention of cortical neuronal degeneration (Vaur *et al*., 2017). Depletion of the enzyme nicotinamide phosphoribosyltransferase (NAMPT), rate-limiting for NAD^+^ biosynthesis, in mouse skeletal muscle severely diminishes NAD^+^ levels and induces sarcopenia. Oral repletion of NAD^+^ with NR in this model rescued pathology in skeletal muscle in a cell-autonomous manner (Frederick *et al*., 2016). However, recent data in mice tracing NAD^+^ fluxes questioned whether oral NR has the ability to access muscle (Liu *et al*., 2018). Thus, whether oral NR can augment the human skeletal muscle NAD^+^ metabolome is currently unknown.

A decline in NAD^+^ availability and signalling appears to occur as part of the aging process in many species (Gomes *et al*., 2013; Mouchiroud *et al*., 2013), though there is a paucity of data to confirm that this is the case in human aging. NR and nicotinamide mononucleotide (NMN) are reported to extend life span (Zhang *et al*., 2016) and enhance metabolism in aged mice (Mills *et al*., 2016). To date, NR supplementation studies in humans have been reported, focussing on cardiovascular (Martens *et al*., 2018), systemic metabolic (Dollerup *et al*., 2018), and safety (Conze *et al*, 2019) end-points, but have not addressed advanced aging, tissue metabolomic changes, or effects on muscle metabolism and function.

Herein, we set out to study if oral NR is available to aged human skeletal muscle and whether potential effects on muscle metabolism can be detected. We conducted a 21-day NR supplementation intervention in a cohort of 70 – 80 year old men in a randomized, double-blind, placebo-controlled crossover trial. We demonstrate that NR augments the skeletal muscle NAD^+^ metabolome inducing a gene expression signature suggestive of downregulation of energy metabolism pathways, but without affecting muscle mitochondrial bioenergetics or metabolism. Additionally, NR suppresses specific circulating inflammatory cytokines levels. In an additional study, we used ^31^P magnetic resonance spectroscopy (MRS) and show that NAD^+^ decline is not associated with chronological aging *per se* in either human muscle or brain.

## RESULTS

### Oral NR is safe and well tolerated in aged adults

Twelve aged (median age 75 years) and marginally overweight (median BMI 26.6 kg/m^2^; range 21 - 30), but otherwise healthy men were recruited and orally supplemented with NR 1 g per day for 21-days in a randomized, double-blind, placebo-controlled crossover design with 21-days washout period between phases. Baseline characteristics of participants are included in Suppl Table 1. NR chloride (Niagen ®) and placebo were provided as 250 mg capsules (ChromaDex, Inc.) and subjects were instructed to take two in the morning and two in the evening. All participants completed the study visits (5 in total) and assessments according to protocol (Suppl Figure 1). Visit 1 was a screening and enrolment visit, while visit 4 was after the washout period and only fasting blood and 24hr urine were collected. The protocol design for visits 2, 3, and 5 included muscle biopsy, fasting blood analyses, glucose tolerance test, muscle arterio-venous difference technique, venous occlusive plethysmography, and indirect calorimetry analysis (Suppl Figure 1). NR was well tolerated and screening for a range of haematological and clinical biochemistry safety parameters (including renal, liver and thyroid functions), revealed no adverse effects (Suppl Table 2). No clinical adverse events were reported during the intervention in either phase. Of note, four participants (33.3%), blinded to the intervention arm, self-reported a noticeable increase in libido whilst on NR. There were no such reports whilst on placebo.

**Figure 1.**
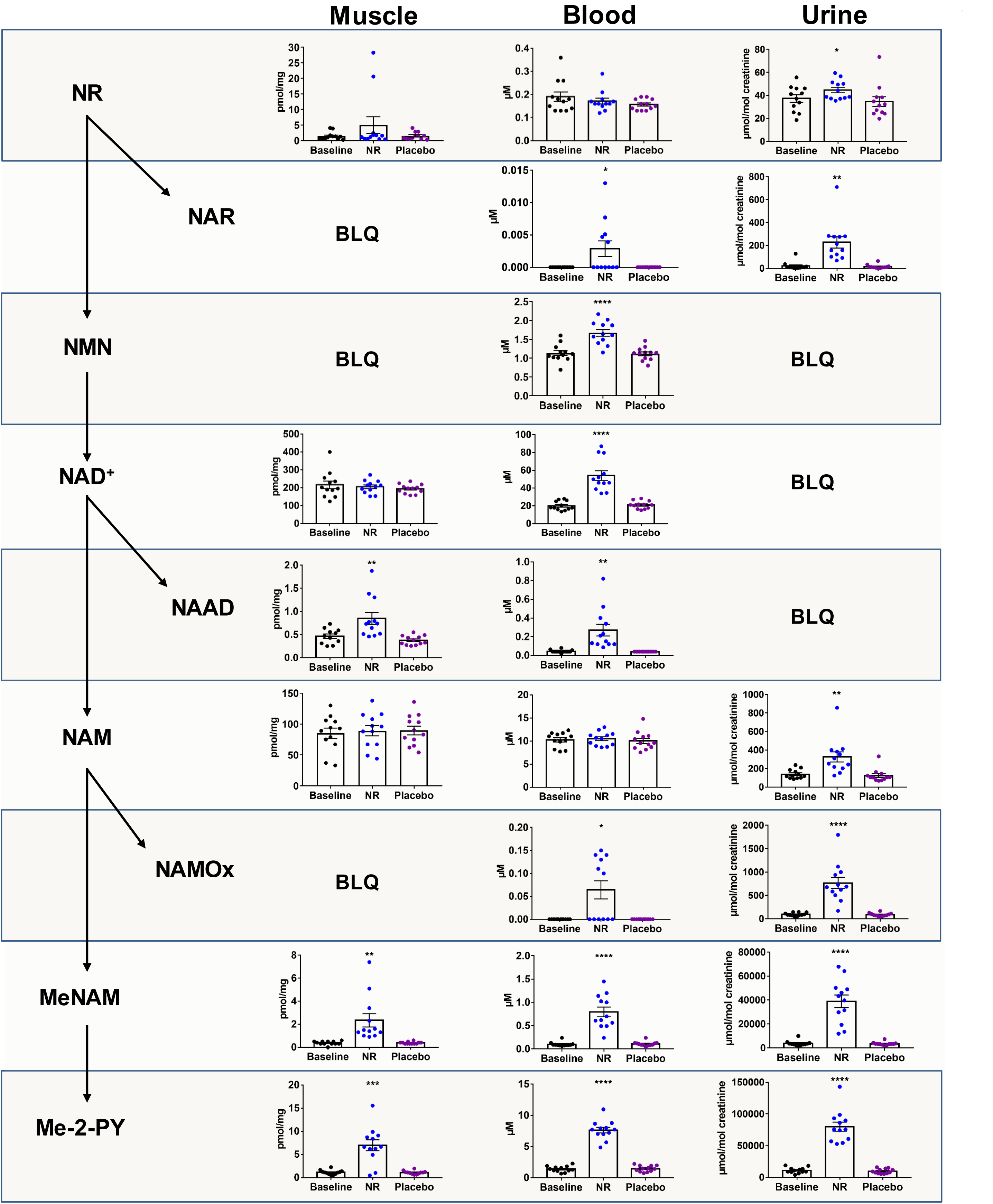
NR augments the human skeletal muscle NAD^+^ metabolome. Schematic representation of nicotinamide riboside (NR) metabolism within the nicotinamide adenine dinucleotide (NAD^+^) metabolome, accompanied by observed levels of metabolites measured using LC-MS/MS in skeletal muscle, whole blood, and urine, at baseline and after each of the NR and placebo periods. NAD^+^ metabolomics data at the end of the washout period are shown In **Suppl Table 3**. Skeletal muscle data were normalized to the weight of muscle pellet used for extraction. Urine data were normalized to urinary creatinine. BLQ, below limit of quantification; NMN, nicotinamide mononucleotide; NAAD, nicotinic acid adenine dinucleotide; NAM, nicotinamide; NAMOx,, nicotinamide N-oxide; MeNAM, N-methyl nicotinamide; Me-2-py, N1-Methyl-2-pyridone-5-carboxamide. Oher metabolites are shown in Suppl. Figure 2. Data are obtained from 12 participants at each phase and presented as mean ± SEM. Significance was set at p < 0.05 using paired t-test, and represents difference between NR and placebo and between NR and baseline. The absence of significance symbols indicates lack of statistical significance.

### Oral NR augments the skeletal muscle NAD^+^ metabolome

To assess the effects of NR supplementation on NAD^+^ metabolism, we used a targeted LC-MS/MS method (Trammell and Brenner, 2013) to quantify the NAD^+^ metabolome in skeletal muscle, whole venous blood, and urine. We examined the NAD^+^ metabolome in skeletal muscle biopsies from all participants in a fasted state at baseline and after the NR and placebo phases, 14 h after the last dose and prior to the physiological assessments. Samples were collected 14 h after the last dose so participants could attend in a fasted state and also to evaluate the effects of longer-term NR administration rather than those of an acute dose. Fourteen metabolites were measured in muscle extracts (Figure 1, Suppl Figure 2A, and Suppl Table 3). NR was detectable in muscle but was not elevated in the NR supplementation period (NR 1.4 pmol/mg µM vs. placebo 1.25 pmol/mg; p = 0.23). Consistent with nicotinic acid adenine dinucleotide (NAAD) as a highly sensitive biomarker of NR supplementation and enhanced rate of NAD^+^ synthesis (Trammell et al. 2016), we found that oral NR resulted in a 2-fold increase in muscle NAAD (NR 0.73 pmol/mg vs. placebo 0.35 pmol/mg; p = 0.004), without an increase in NAD^+^ (NR 210 pmol/mg vs. 197 pmol/mg; p = 0.22). NR supplementation did not affect muscle nicotinamide (NAM) (NR 92.0 pmol/mg vs. placebo 86.5 pmol/mng; p = 0.96). However remarkably, we detected 5-fold increases in the products of NAM methylation clearance pathways; N-methyl nicotinamide (MeNAM; NR 1.45 pmol/mg vs. placebo 0.35 pmol/mg; p = 0.006), N1-Methyl-2-pyridone-5-carboxamide (Me-2-py; NR 6.6 pmol/mg vs. placebo 1.1 pmol/mg; p = < 0.001), and N1-Methyl-4-pyridone-5-carboxamide (Me-4-py; NR 1.6 pmol/mg vs. placebo 0.3 pmol/mg; p = < 0.001) (Figure 1, Suppl Figure 2A, **and Suppl Table 3**).

**Figure 2.**
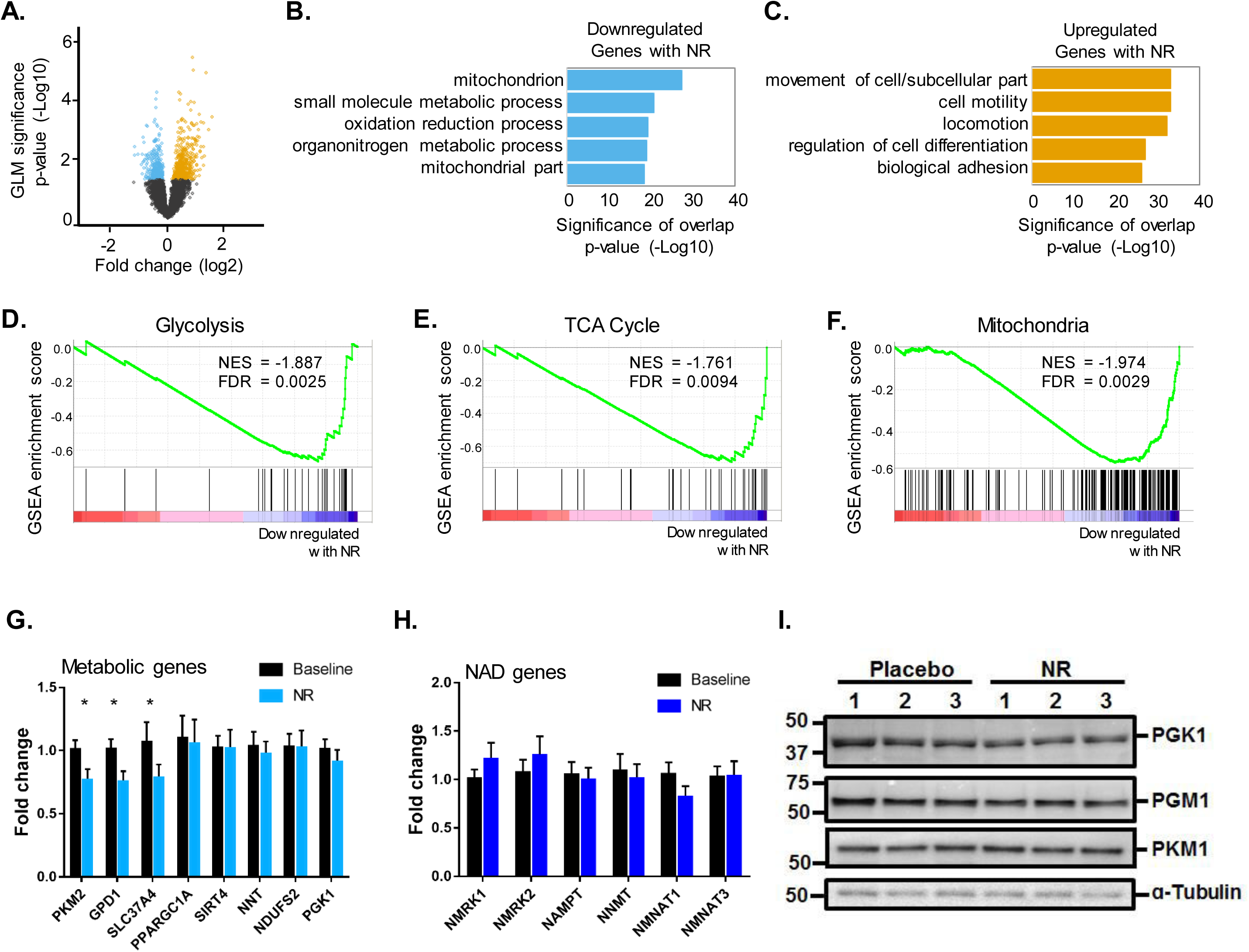
NR supplementation induces a transcriptional signature in human skeletal muscle. (A) Differential gene expression analysis on baseline and nicotinamide riboside (NR) treated muscle samples (n=12 at each phase). Volcano plot of differential gene expression between baseline and NR treated human muscle samples. Fold change (Log2, x-axis) of gene expression is plotted against p value for differential gene expression (-Log10, y-axis). Coloured dots represent Ensembl genes that are either upregulated (in orange) or downregulated (in blue) upon NR supplementation at a p value < 0.05. (B) and (C) Gene ontology analysis of significantly dysregulated genes upon NR supplementation where (B) is downregulated genes and (C) is upregulated genes. Gene ontology analysis was performed using GSEA. Bars represent the p value (-Log10) of overlap from hypergeometric distribution. (D) Gene set enrichment analysis (GSEA) suggests that genes belonging to the gene set “Glycolysis” are downregulated upon NR supplementation. Normalized enrichment score (NES) and nominal p-value is presented on the top left corner of the graph. (E) As in (D) for genes involved in the TCA cycle (F) As in (D) for genes involved in the gene set “mitochondria” (G) Quantitative PCR analysis of a select panel of downregulated genes identified through differential gene expression analysis. GAPDH was used as housekeeping gene. Error bars represent SEM (n=12). (H) As in (G) for NAD^+^ pathway related genes. (I) Quantification of phosphoglycerate kinase 1 (PGK1), phosphoglucomutase 1 (PGM1), and pyruvate kinase M1 (PKM1) proteins using immunoblotting assay. Tubulin was used as a loading control. Data are obtained from 12 participants at each phase and wherever relevant are presented as mean ± SEM. Significance was set at p < 0.05. The absence of significance symbols indicates lack of statistical significance.

In the blood, we measured 15 metabolites from each participant at baseline and following each of the NR, placebo, and washout periods (Figure 1, Suppl Figure 2B, **and Suppl Table 3**). NR was also detectable in the blood but was not increased compared to placebo at 14 h after the last dose of NR (NR 0.16 µM vs. placebo 0.15 µM; p = 0.31). This is expected as the predicated Cmax for NR is approximately 3 h (Airhart *et al*., 2017). NR increased the concentrations of NAD^+^ > 2-fold (NR 47.75 µM vs. placebo 20.90 µM; p = <0.001) and NMN 1.4-fold (NR 1.63 µM vs. placebo 1.13 µM; p = <0.001). A recent study reported that oral NR is rapidly metabolised in the liver to NAM, which can enhance tissue NAD^+^ metabolomes (Liu *et al*., 2018). However, chronic NR supplementation did not elevate NAM in the blood (NR 10.60 µM vs. placebo 9.50 µM; p = 0.41). Again, NAM urinary clearance pathways were highly active following NR with marked excess of MeNAM (NR 0.66 µM vs. placebo 0.10 µM; p = <0.001), Me-2-py (NR 7.69 µM vs. placebo 1.44 µM; p = <0.001), and Me-4-py (NR 3.82 µM vs. placebo 0.48 µM; p = <0.001) (Figure 1, Suppl Figure 2B, **and Suppl Table 3**). NR elevated blood NAAD levels by 4.5-fold (NR 0.18 µM vs. Placebo 0.04 µM; p = <0.001).

Urinary NAD^+^ metabolomics showed that NR was detectable and increased with NR supplementation (NR 41.5 µmol/mol creatinine vs. Placebo 31.7 µmol/mol creatinine; p = 0.02) (Figure 1). Furthermore, a near 20-fold increase in nicotinic acid riboside (NAR; NR-185.5 µmol/mol creatinine vs. placebo-10.3 µmol/mol creatinine; p = 0.001) was observed.

This observation may support the suggestion that NR supplementation leads to retrograde production of NAAD, nicotinic acid mononucleotide (NAMN), and NAR (Trammell et al. 2016). However, direct NR transformation into NAR cannot be excluded. Unlike muscle and blood, NAM was elevated in the urine 2.5-fold (NR-282 µmol/mol creatinine vs. Placebo-106.5 µmol/mol creatinine; p = 0.004). These data establish the extent and breadth of changes to NAD^+^ metabolites in human muscle, blood, and urine after NR supplementation. The data indicate that oral NR greatly boosts the blood NAD^+^ metabolome without an increase in NAM, increases muscle NAD^+^ metabolism, and leads to disposal of urinary clearance products.

### Oral NR results in downregulation of gene sets associated with energy metabolism in skeletal muscle

We next assessed NR-mediated transcriptional changes in skeletal muscle. RNA sequencing followed by differential gene expression (DGE) analysis of muscle biopsies from the 12 participants revealed 690 upregulated and 398 downregulated genes between baseline and NR supplementation at p value < 0.05 (Figure 2A **and Suppl Table 4**). Using gene annotation analysis (DAVID and GSEA) (Mootha *et al*., 2003; Subramanian *et al*., 2005; Huang, Brad T Sherman and Lempicki, 2009; Huang, Brad T. Sherman and Lempicki, 2009), we examined the enrichment of genes that belong to known molecular pathways in our list of up- or downregulated genes. Our results suggest that genes significantly downregulated with NR supplementation were enriched in pathways relating to energy metabolism including those of glycolysis, TCA cycle and mitochondria (Figure 2B, and Suppl Table 5). This is consistent with the recent discovery that oral NR depresses mitochondrial membrane potential while improving blood stem cell production in mice (Vannini *et al*., 2019).

Pathways upregulated upon NR supplementation prominently belonged to gene ontology categories such as cell adhesion, actin cytoskeleton organization, and cell motility (Figure 2C). This supports a previously identified role for the NAD^+^-generating enzyme nicotinamide riboside kinase 2b (Nrk2b) in zebrafish skeletal muscle cell adhesion (Goody *et al*., 2010).

We next examined all the genes that belonged to the glycolysis, mitochondrial and the TCA cycle pathways and found that they were predominantly downregulated following NR supplementation, whereas 10 control genes sets, of the same size and expression level were not (Figure 2D-F and Suppl Figure 3A). Similarly, we found that the genes belonging to the gene ontology terms actin filament-based process, cell motility, and biological cell adhesion, were mainly upregulated upon NR supplementation (Suppl Figure 3B **and** C).

**Figure 3.**
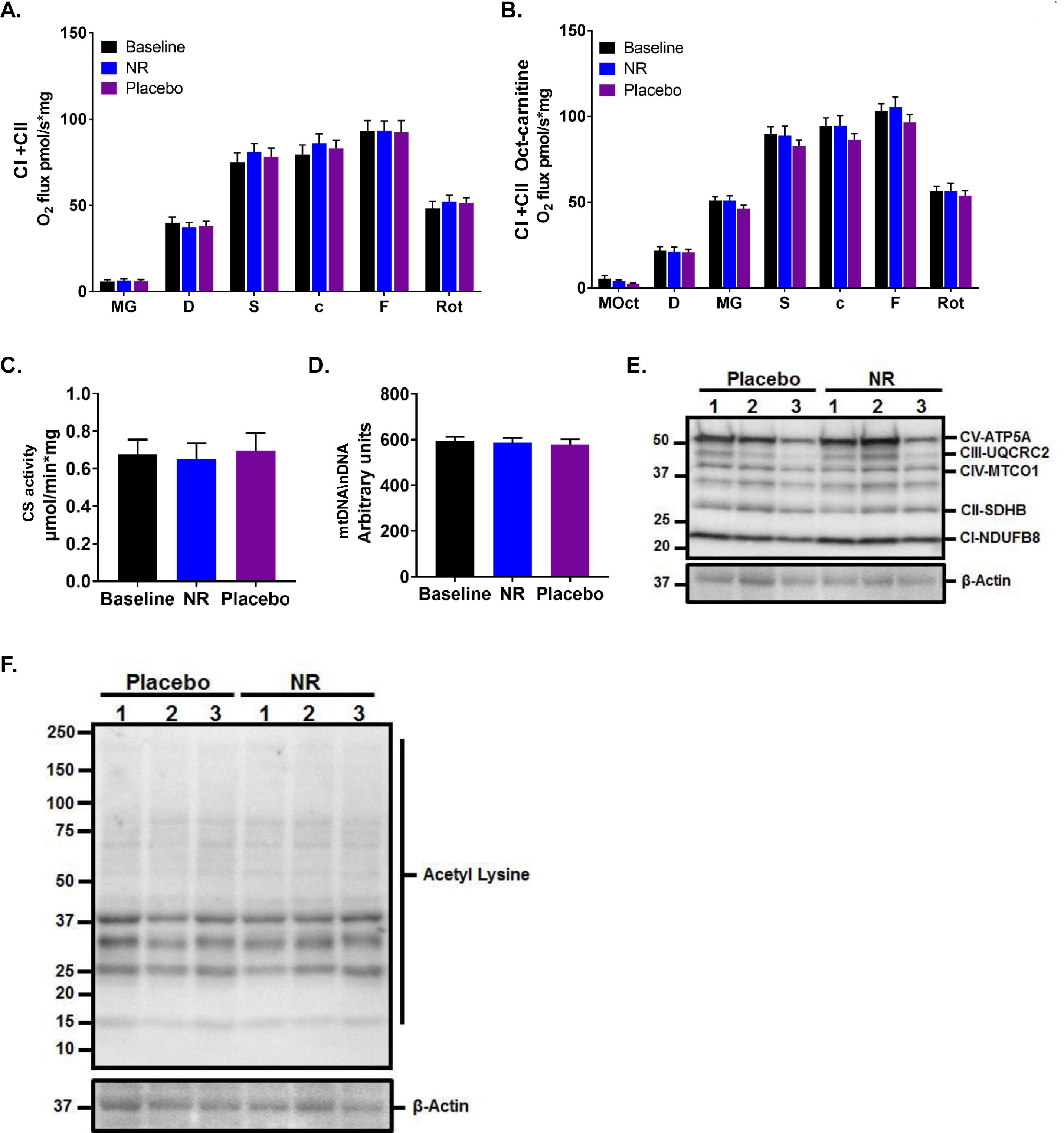
Human skeletal muscle mitochondrial bioenergetics remain unaltered with NR supplementation. (A) Mitochondrial respiration of permeabilized muscle fibres upon the addition of Complex I and Complex II substrates at baseline and after 3 weeks supplementation of NR and placebo. MG, malate and glutamate; D, ADP; S, succinate; c, cytochrome C; F, FCCP; Rot, rotenone. Data normalised to muscle fibre weight. (B) Mitochondrial respiration as per (A) but with the prior addition of the fatty acid conjugate octanoyl-carnitine in addition to malate (MOct). (C) Citrate synthase (CS) activity in human skeletal muscle at baseline and after NR and placebo. (D) Relative PCR expression of mitochondrial DNA (mtDNA) to nuclear DNA (nDNA) at baseline and after NR and placebo, expressed as arbitrary units. (E) Western blot showing the expression of selected mitochondrial proteins in skeletal muscle lysates compared to β-Actin as housekeeping protein. (F) Western blot showing the expression of acetylation proteins in skeletal muscle lysates compared to β-Actin as housekeeping protein. Data are obtained from 12 participants at each phase and wherever relevant are presented as mean ± SEM. Significance was set at p < 0.05. The absence of significance symbols indicates lack of statistical significance.

In agreement with DGE analysis, quantitative real time PCR showed downregulation of selected genes involved in energy metabolism (Figure 2G). We found no changes in the transcript levels of key genes involved in NAD^+^ metabolism, corroborating the DGE analysis (Figure 2H). We also verified some of the upregulated targets by quantitative PCR (Suppl Figure 3D), and undertook some immunoblotting validation (Suppl Figure 3D).

As it has previously been shown that NR increases glycolysis in mouse cardiac cells (Diguet *et al*., 2018), and because our data do not support an NR-mediated transcriptional upregulation of glycolysis related genes, we examined protein expression levels of glycolytic enzymes in our muscle biopsies and show them to be unchanged after NR (Figure 2I).

### Three weeks of oral NR does not alter skeletal muscle mitochondrial bioenergetics or hand-grip strength

Several preclinical studies suggest that NR enhances mitochondrial energy programmes in skeletal muscle (Cantó *et al*., 2012; Frederick *et al*., 2016) through mechanisms that involve redox and sirtuins activation. Therefore, we undertook detailed assessment of muscle mitochondrial respiration in biopsies after NR supplementation using high-resolution respirometry, the gold standard method for the *ex vivo* assessment of mitochondrial function. No differences were detected between NR and placebo groups in skeletal muscle complex I- and complex II-mediated oxidative phosphorylation and maximal respiratory capacity, with (Figure 3A) and without the prior addition of the fatty acid conjugate octanoyl-carnitine (Figure 3B). In line with this, the activity of citrate synthase, commonly used as a quantitative measure of mitochondrial content (Larsen *et al*., 2012) (Figure 3C), and mitochondrial copy number (mtDNA) (Phielix *et al*., 2008) (Figure 3D) were unchanged by NR supplementation. Similarly, levels of skeletal muscle biopsy mitochondrial resident proteins, directly involved in the electron transport chain, were unaltered upon NR supplementation (Figure 3E). We then tested whether the NR-driven increase in the NAD^+^ metabolome translates into higher sirtuin-mediated deacetylation activity and performed western blotting to assess pan-acetylation status, but again did not detect NR-mediated changes to muscle protein acetylation (Figure 3F).

Data from rodents suggest that NAD^+^ supplementation can improve physiological function in skeletal muscle decline (Cantó *et al*., 2012; Frederick *et al*., 2016; Mills *et al*., 2016), thus we used hand-grip strength as a surrogate marker for muscle function. Hand-grip strength correlates with leg strength, and is used for the diagnosis of sarcopenia and frailty, and is a better predictor for clinical outcomes than low muscle mass (Lauretani *et al*., 2003). A decline in hand-grip strength is observed after the third decade of life (when median peak strength is 51 kg of force in men) (Dodds *et al*., 2014), dropping to median of 33.8 kg of force in our participants. A grip strength of <30 kg of force in men is a diagnostic criterion for sarcopenia (Cruz-Jentoft *et al*., 2010). After 3 weeks of supplementation, we did not observe any differences in the participants’ peak hand-grip strength (NR 32.5 kg vs. placebo 34.7 kg; p = 0.96) or body weight adjusted relative strength (NR 2.4 vs. placebo 2.3; p = 0.96) between NR and placebo (Suppl Figure 4).

**Figure 4.**
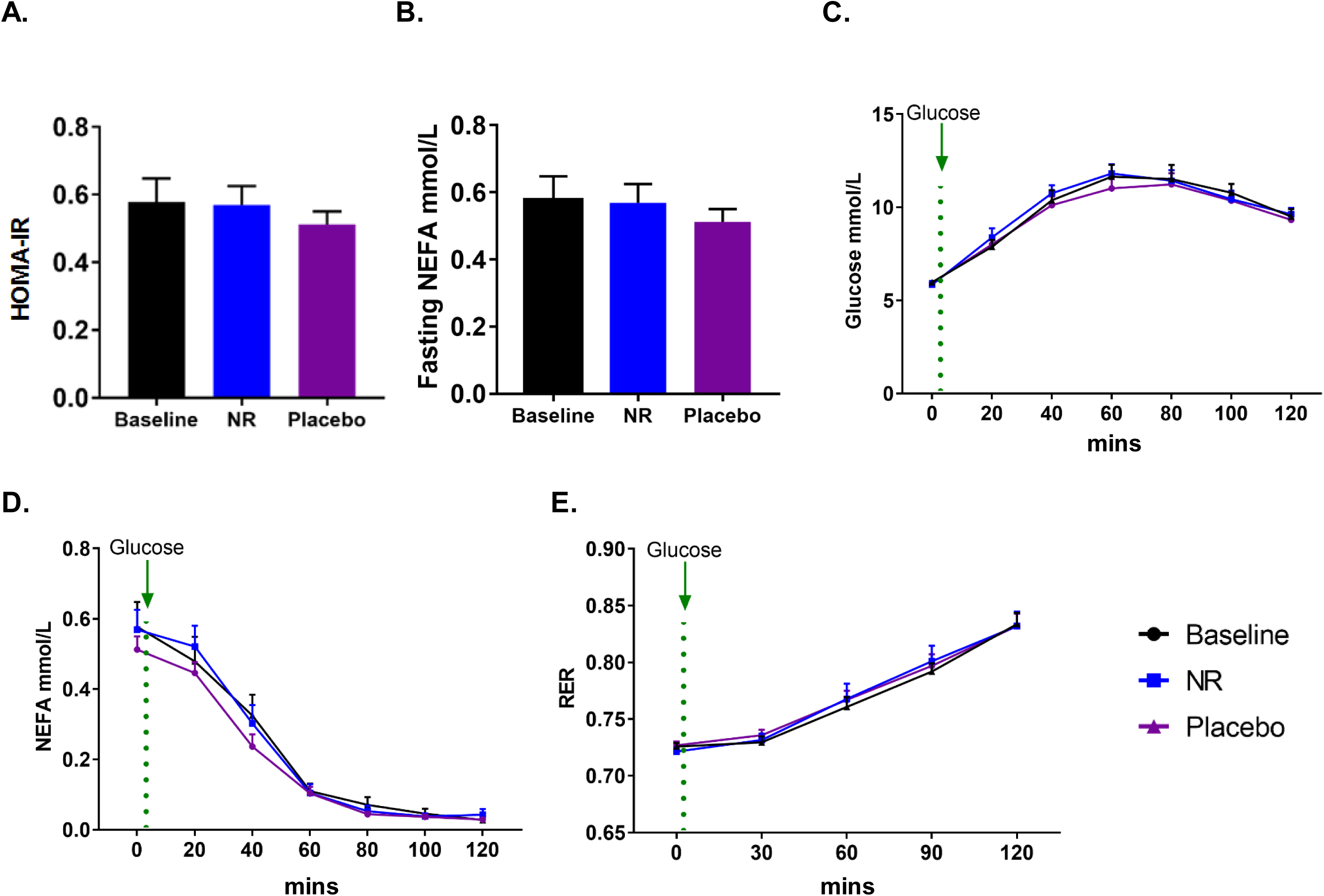
Systemic readouts of metabolism are unaltered with NR supplementation. (A) HOMA-IR at baseline and after nicotinamide riboside (NR) and placebo. (B) Fasting non-esterified fatty acid (NEFA) level at baseline and after NR and placebo. (C) Plasma glucose response in a glucose tolerance test at baseline and after NR and placebo. The green dotted line represents when 75g oral glucose load was taken. (D) Plasma NEFA response in a glucose tolerance test at baseline and after NR and placebo. The green dotted line represents when 75g oral glucose load was taken. (E) Respiratory exchange ratio (RER) at baseline and after NR and placebo. The green dotted line indicates when 75g oral glucose was taken. Data are obtained from 12 participants at each phase and presented as mean ± SEM. Significance was set at p < 0.05. The absence of significance symbols indicates lack of statistical significance.

### Oral NR does not alter systemic cardiometabolic parameters

Several preclinical studies have described that NAD^+^ supplementation promotes resistance to weight gain, ameliorates markers of cardiometabolic risk, and improves metabolic flexibility (Yoshino, Baur and Imai, 2017). As NR increased the circulating levels of the NAD^+^ metabolome, we reasoned that there was increased NAD^+^ availability and turnover in central and peripheral tissues and assessed for resultant cardiometabolic adaptations. Two studies—one of 12 weeks of NR supplementation at 2 g/day) in subjects with obesity (Dollerup *et al*., 2018) and one of 6 weeks of NR supplementation at 1 g/day in older adults (Martens *et al*., 2018) suggested potential benefits with respect to fatty liver and blood pressure, respectively. Data for participants at baseline and following NR or placebo are reported in Suppl Table 1. There were no changes in body weight, blood pressure, lipid profile, fasting glucose and insulin (Suppl Table 1), and HOMA-IR (Figure 4A). A rebound increase in non-esterified fatty acids (NEFA) has previously been associated with the nicotinic acid analog, acipimox (van de Weijer *et al*., 2015), however, NR did not produce this effect in our trial (Figure 4B). Glucose handling was studied using an oral glucose tolerance test with no effect of NR measured in glucose levels during the 2-hour test (Figure 4C). Following the oral glucose load and the consequent insulin stimulation, NEFA levels were appropriately suppressed and no difference in this response was observed between NR and placebo (Figure 4D). We also assessed metabolic flexibility using indirect calorimetry to derive respiratory exchange ratios (RER; calculated as VCO_2_ expired/ VO_2_ consumed) reflecting whole body metabolic substrate use. Measurements were initiated in the fasted state and monitored during the response to the oral glucose load. Median fasting RER was appropriate at 0.72 and 0.73 for the NR and placebo periods, respectively (p = 0.68). In response to glucose, RER values significantly increased indicating adequate switching from lipids towards carbohydrates utilisation, with no differences in response to 3 weeks of NR supplementation observed at 2h (RER 0.83 and 0.84 for NR and placebo, respectively) (Figure 4E).

### Oral NR does not alter skeletal muscle blood flow or substrate utilisation

Recent mouse data showed that NR increases angiogenesis and muscle blood flow (Das *et al*., 2018). Therefore, we used venous occlusive plethysmography to test forearm muscle blood flow in the participants in a non-invasive manner (Greenfield, Whitney and Mowbray, 1963). At fasting, no NR-mediated differences were detected in muscle blood. Following oral glucose load, muscle blood flow gradually increases, but again with no differences between NR and placebo (Figure 5A).

**Figure 5.**
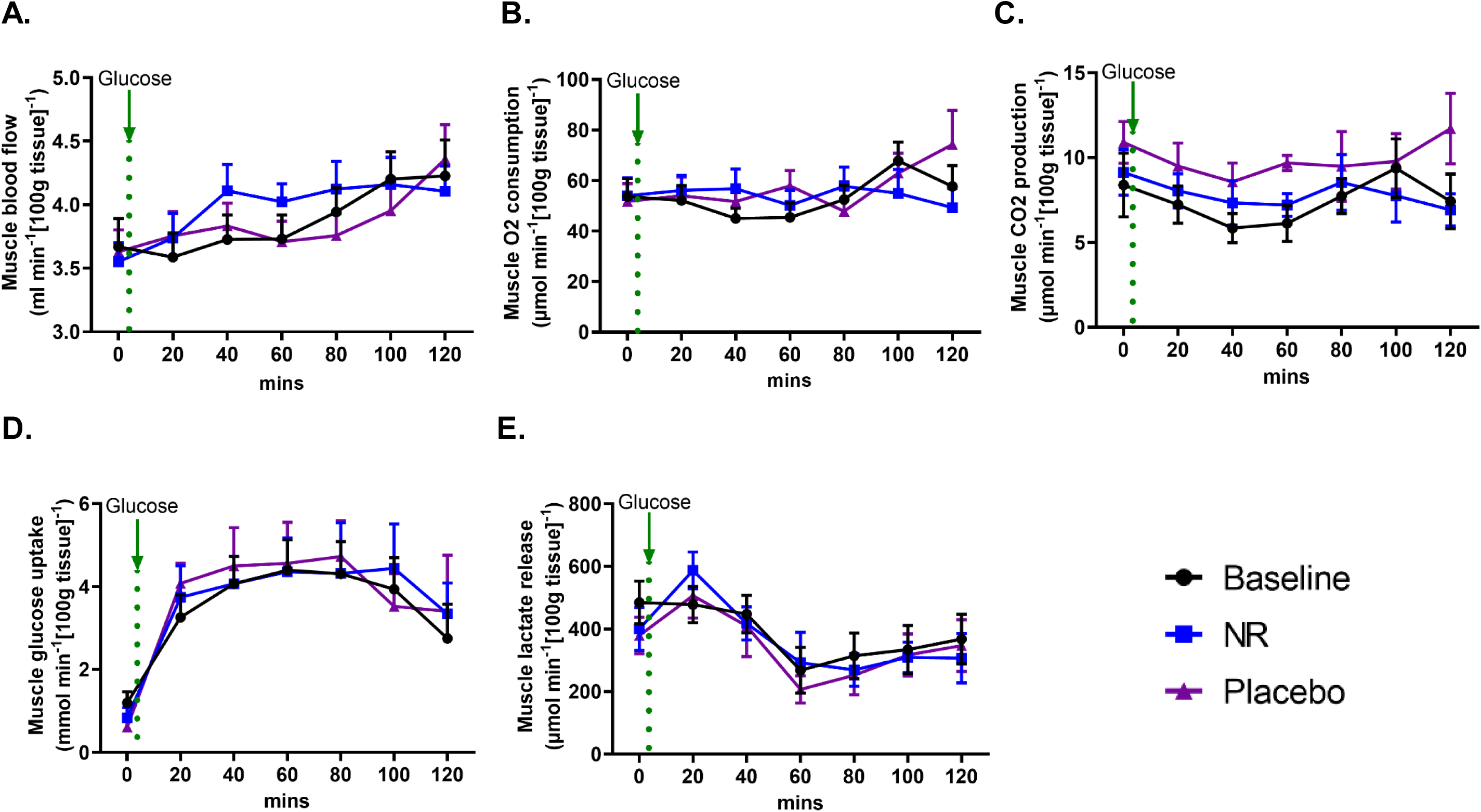
Forearm muscle blood flow and substrate utilisation are unaffected by NR supplementation. (A) Muscle blood flow using venous occlusive plethysmography at baseline and after the nicotinamide riboside (NR) and placebo phases. The green dotted line represents when 75g oral glucose load was taken. (B) Muscle O_2_ consumption and (C) CO_2_ production at baseline and after NR and placebo. The green dotted line represents when 75g oral glucose load was taken. (D) Muscle glucose uptake and (E) Lactate release at baseline and after NR and placebo. The green dotted line represents when 75g oral glucose load was taken. Data are obtained from 12 participants at each phase and presented as mean ± SEM. Significance was set at p < 0.05 using paired t-test. The absence of significance symbols indicates lack of statistical significance.

We then used the arteriovenous difference method (**see methods section**) to compare substrate utilisation across the forearm muscle (between arterial blood supplying the muscle and venous blood drained from the muscle), with muscle blood flow taken into consideration (Bickerton *et al*., 2007). No differences were detected in O_2_ consumption (Figure 5B) and CO_2_ production (Figure 5C) between NR and placebo at the fasting state and in response to oral glucose. Muscle glucose uptake was increased following oral glucose before gradual decline. Similar to blood glucose, no changes were observed in muscle glucose handling with or without NR (Figure 5C). Oral glucose reduced lactate production from muscle, again without a difference in response between NR and placebo (Figure 5E). These data are concordant with our observations on systemic readouts of metabolism, and suggest that the skeletal muscle transcriptomic signature of downregulated mitochondrial and glycolysis genes is undetectable when considered at a functional level.

### Oral NR depresses circulating levels of inflammatory cytokines

Chronic inflammation appears to be a consistent feature of aging, even in apparently healthy individuals (Singh and Newman, 2011), and may contribute to age-related disturbance in metabolic homeostasis (Imai and Yoshino, 2013). We hypothesized that NR supplementation would reduce the levels of circulating inflammatory cytokines. We measured multiple inflammatory cytokines (**see methods section for details**), 10 of which were within the assay detection range (Figure 6). NR significantly decreased the levels of the interleukins IL-6 (Figure 6A), IL-5 (Figure 6B), and IL-2 (Figure 6C), and tumor necrosis factor-alpha (TNF-a) (Figure 6D) compared to baseline. We detected a statistically significant difference in the levels of IL-2 between baseline and placebo (Figure 6C), and lack of a difference in levels of TNF-a between NR and placebo despite a difference between NR and baseline (Figure 6D). This is seemingly due to NR carry-over effect beyond the washout period, as evident by period effect analysis **(Suppl** Figure 5A-D), confirming that the cohort randomized to placebo first had no difference in IL-2 between baseline and placebo **(Suppl** Figure 5C) and there was a difference in TNF-a between NR and placebo **(Suppl** Figure 5D). No NR-mediated changes were detected in the serum levels of IL-12 (Figure 6E), IL-8 (Figure 6F), interferon-gamma (IFN-g) (Figure 6G), monocyte chemoattractant protein-1 (MCP-1) (Figure 6H), macrophage inflammatory protein-1 beta (MIP-1B) (Figure 6I), and high-sensitivity C-reactive protein (hsCRP) (Figure 6J). Thus, it will be interesting to further investigate depressed IL-6, IL-5, IL-2 and TNF-a as biomarkers and/or mediators of oral NR in rodent models and humans.

**Figure 6.**
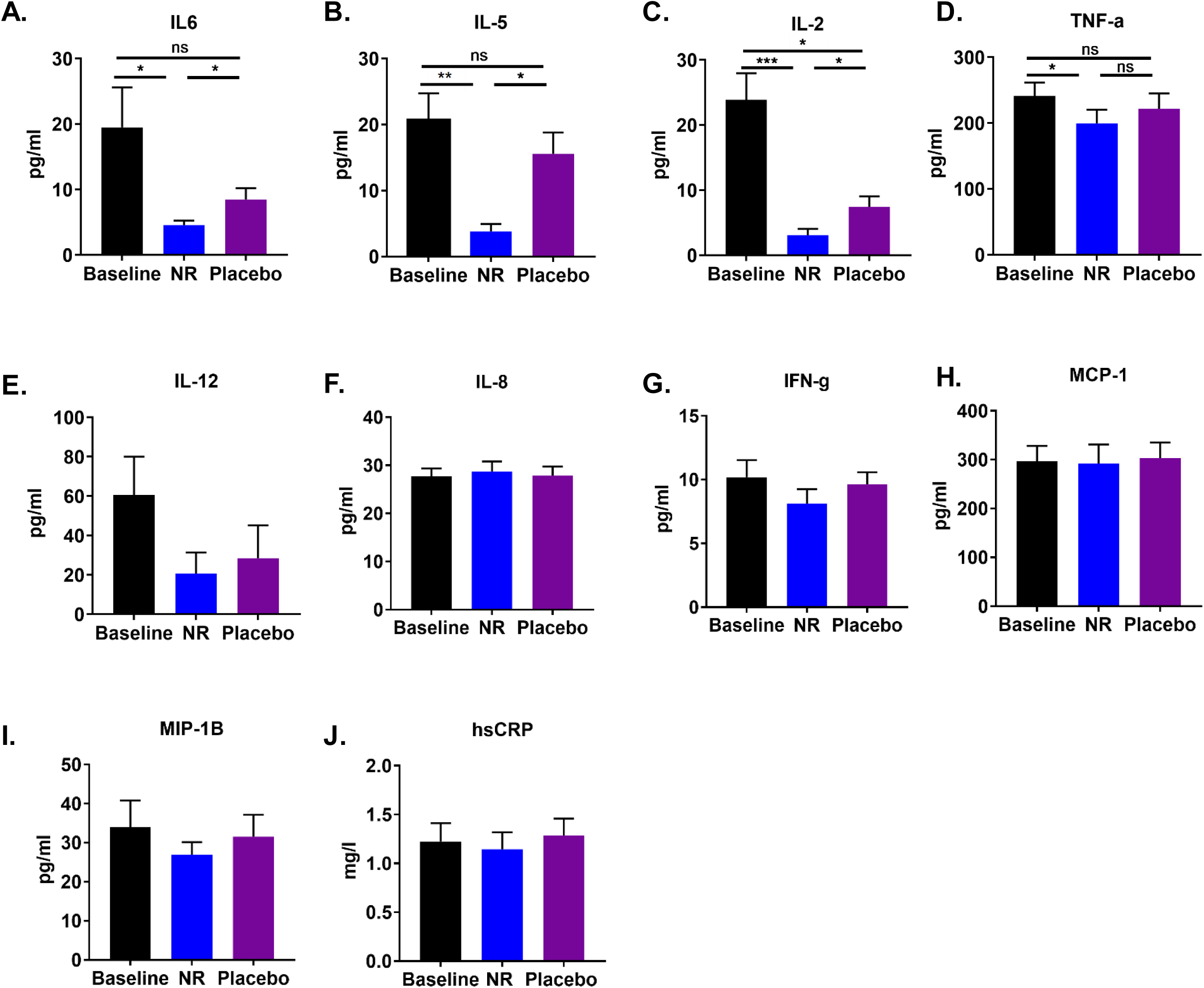
NR supplementation suppresses the circulating levels of inflammatory cytokines. Levels of serum inflammatory cytokines at baseline and after each of the nicotinamide riboside (NR) and placebo phases, including: (A) interleukin 6 (IL-6), (B) interleukin 5 (IL-5), (C) interleukin 2 (IL-2), (D) tumor necrosis factor-alpha (TNF-a), (E) interleukin 12 (IL-12), (F) interleukin 8 (IL-8), (G) interferon-gamma (IFN-g), (H) monocyte chemoattractant protein-1 (MCP-1), (I) macrophage inflammatory protein-1 beta (MIP-1B), and (J) high-sensitivity C-reactive protein (hsCRP). Data are obtained from 12 participants at each phase and presented as mean ± SEM. Significance was set at p < 0.05 using paired t-test. The absence of significance symbols indicates lack of statistical significance.

### NAD^+^ content in skeletal muscle and brain does not decline with age

We questioned whether there was any degree of pre-existing NAD^+^ deficiency in aged humans as previously described in laboratory mice (Gomes *et al*., 2013; Mouchiroud *et al*., 2013). To address this, we investigated whether tissue NAD^+^ content is different between healthy young and aged individuals. We recruited a young cohort (n=16, median age 21 years ± 4, male:female 5:11, mean BMI 22.9 kg/m^2^ ± 2.9) and an aged cohort (n=11, median age 69 years ± 4, male:female 5:6, mean BMI 23 kg/m^2^ ± 2.3) and used ^31^P magnetic resonance spectroscopy (MRS) to measure calf muscle NAD^+^ content *in vivo*. We did not detect significant differences between the age groups (Figure 7A). It was not possible to reliably differentiate the magnetic resonance signal of NAD^+^ from that of NADH in the calf muscle due to reduced magnetic homogeneity, resulting in reduced spectral quality (increased linewidth). We also compared brain NAD(H) content using ^31^P MRS in young (n=12, mean age 21± 4, male:female 4:8, mean BMI 22.9 kg/m^2^ ± 2.9) and aged individuals (n=11, mean age 69 years ± 4, male:female 4:7, mean BMI 23 kg/m^2^ ± 2.3) and again found no significant differences between the two groups in the concentrations of NAD^+^, NADH, and NAD^+^ redox potential (Figure 7B**-F**). Previous work using ^31^P MRS reported an age-dependent decline in human brain NAD^+^ concentration and redox potential (Zhu *et al*., 2015), although between age-group comparisons were not formally tested in that study – presumably due to small sample sizes (n=7, 4, 6 for young, middle-aged and older groups, respectively). Nevertheless, their values of brain NAD^+^ concentration and redox potential closely resemble our data.

**Figure 7.**
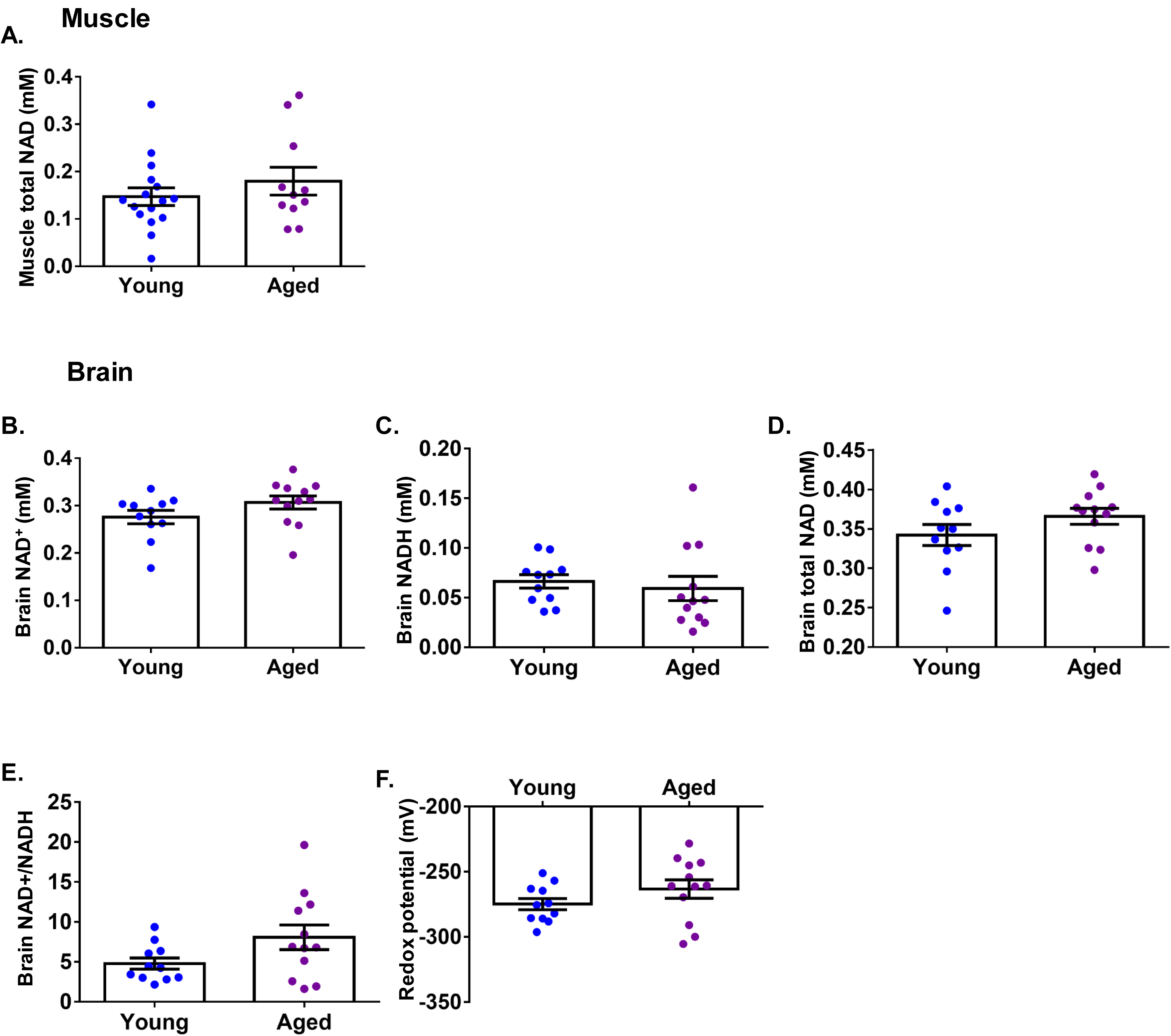
NAD^+^ content in skeletal muscle and brain is not different between young and aged humans. (A) ^31^P magnetic resonance spectroscopy (MRS)-based total NAD^+^ measurement in the calf muscle comparing young (n=16) and aged (n=11) individuals. ^31^P MRS-based measurement of brain (B) NAD^+^, (C) NADH, (D) total NAD^+^, (E) NAD^+^/NADH ratio, and (F) NAD redox potential comparing young (n=12) and aged (n=11) individuals. Data are presented as mean ± SEM. Significance was set at p < 0.05 using independent t-test. The absence of significance symbols indicates lack of statistical significance.

## Discussion

The NAD^+^ precursor NR has been studied extensively in animal and cell models. Its application *in vivo* has produced impressive results ameliorating metabolic dysfunction and muscle decline (Fang *et al*., 2017). No data exist on whether oral NR is available to human skeletal muscle, and data on tissue NAD^+^ content during aging are sparse, as are the consequences of NR supplementation in aged humans. We conducted two human clinical studies to address these issues.

Using a robust clinical trial design, we show that 21-days of NR supplementation is safe and well tolerated in an aged male cohort and leads to an enhanced NAD^+^ metabolome in whole blood, corroborating data recently reported by others (Martens et al. 2018; Trammell et al. 2016). The median BMI in this trial is 26.6 kg/m^2^ (i.e. slightly overweight), but this is highly prevalent in aged populations (Winter *et al*., 2014), and may not indicate unhealthy state (Porter Starr and Bales, 2015).

Experiments in genetic mouse models have shown that oral NR is available to cardiac (Diguet *et al*., 2018) and skeletal (Frederick *et al*., 2016) muscle, though it was also suggested that the benefit of extrahepatic NAD^+^ from oral NR is mediated by circulating NAM (Liu *et al*., 2018). Here we show that oral NR increased human skeletal muscle NAAD, which was previously reported as a more sensitive marker of increased NAD^+^ metabolism than NAD^+^ per se (S. A. Trammell *et al*., 2016), as well as MeNAM, Me-4-py, and Me-2-pywithout a rise in circulating NAM. Previous preclinical studies have established that oral NR is able to functionally restore muscle NAD^+^ despite loss of NAM salvage (Frederick *et al*., 2016; Diguet *et al*., 2018). Although it is clear in rodent models that NR requires NR kinase actvity in muscle (Ratajczak *et al*., 2016; Fletcher *et al*., 2017), further stuides are required to undersand NR dynamics in human muscle cells and tissues. Increased circulating levels of MeNAM and expression of its generating enzyme nicotinamide-N-methyltransferase (NNMT) have been associated with insulin resistance and type 2 diabetes (Kannt *et al*., 2015; Liu *et al*., 2015). However, the NR-mediated abundance of Me-NAM did not alter glucose tolerance or substrate utilization in our study.

The levels of elevated NAM excretory products in skeletal muscle may be a result of pre-existing NAD^+^ sufficiency in this aged cohort and may explain the lack of effect on mitochondrial, physiological, and cardiometabolic parameters. Indeed, use of ^31^P MRS-based skeletal muscle NAD^+^ measurement demonstrated that NAD^+^ levels are similar between young and aged cohorts. A limited number of studies have reported minor age-related declines in NAD^+^ in human tissues (Massudi *et al*., 2012; Chaleckis *et al*., 2016; Zhou *et al*., 2016; Clement *et al*., 2018). It is likely that a “second-hit” arises during chronological aging that leads to tissue NAD^+^ decline and predisposes to age-related disease and frailty. This “second-hit” may be conditions of metabolic stresses such as physical inactivity, chronic inflammation, or presence of a pre-existing cardiometabolic disease (e.g. obesity), and may implicate downregulated NAMPT (Costford *et al*., 2010; Imai and Yoshino, 2013), depressed hepatic NADP(H) (Trammell et al. 2016), and/or activation of CD38 (Camacho-Pereira *et al*., 2016; Covarrubias *et al*., 2019).

We note median hand-grip strength in our participants of 33.8 kg of force, consistent with muscle aging for men in their 8^th^ decade, and likely associated impairment in mitochondrial function. Our data suggest that 3 weeks of NR supplementation without concomitant muscle training is insufficient for increased strength.

NAD^+^ supplementation studies in rodents showed positive effects on muscle structural proteins (Frederick *et al*., 2016; Ryu *et al*., 2016; Zhang *et al*., 2016). It has been suggested that for muscle cell membranes there is capacity for NAD^+^-mediated ADP-ribosylation of integrin receptors that augment integrin and laminin binding and mobilizes paxillin to bind adhesion complexes (Goody *et al*., 2010; Goody and Henry, 2018). Differential gene expression analysis may support this link and may highlight a potential role for NAD^+^ in the maintenance of skeletal muscle architecture, although this NR-induce transcriptomic signature appear to have no functional consequences at protein level after the 21-day supplementation period. This observation may be important as we consider defective integrin and laminin structures such as in the context of muscular dystrophies (Mayer, 2003; McNally, 2012). Our data suggest downregulation of gene sets associated with glycolysis and mitochondrial function, yet our measures of mitochondrial respiration, citrate synthase activity, and mitochondrial copy number were unaltered. Again, expression levels of glycolysis and mitochondrial protein were unchanged with NR in this study. The downregulation of energy-generating processes may be reminiscent of mechanisms associated with calorie restriction (Hagopian, Ramsey and Weindruch, 2003; Ingram and Roth, 2011; Lin *et al*., 2015) or increased mitochondrial quality control as has been observed in blood stem cells (Vannini *et al*., 2019), or suggest that NR can ‘tune’ the expression of energy metabolism pathways to permit a more efficient and potentially stress resilient mitochondrial environment.

Some preclinical studies have reported that NR reduced macrophage infiltration in damaged muscle (Ryu *et al*., 2016; Zhang *et al*., 2016) and attenuated plasma TNF-alpha in models of fatty liver disease (Gariani *et al*., 2016). For the first time in humans, we show significant suppression of a number of circulating inflammatory cytokines. Of note, expression of the NAD^+^-consuming enzyme CD38 increases in inflammatory cells with inflammation (Amici *et al*., 2018), as well as in the blood of aged humans (Polzonetti *et al*., 2012). Supplementing NAD^+^ in this context may be a mechanism mediating the NR-induced anti-inflammatory effects. Though chronic inflammation is a hallmark feature of aging (Singh and Newman, 2011), use of NR may yet find utility in other chronic inflammatory disorders such as chronic obstructive pulmonary disease or rheumatoid arthritis and is worthy of further investigation.

### Overall Conclusions and Limitations of the Study

We report that oral NR augments the aged human skeletal muscle NAD^+^ metabolome while inducing a novel transcriptional signature without affecting mitochondrial function or systemic cardiometabolic parameters. The targeted NAD^+^ metabolome analysis suggests pre-existing NAD^+^ sufficiency, despite hand-grip strength consistent with muscle aging, and this was supported by finding that NAD^+^ content measured by ^31^P MRS in the muscle and brain of a healthy aged cohort was not different from a young cohort. Our data may suggest that chronological age *per se* is not a major factor in altering muscle and brain NAD^+^ metabolism, unlike aged, laboratory mice. A possible limitation of the trial may be the number of participants or the duration of NR administration. However, the sample size was sufficient to detect NR-driven changes in NAD^+^ metabolome, muscle transcriptional signature, and inflammatory profile. Additionally, the transcriptional downregulation of mitochondrial gene sets argues against the lack of bioenergetic NR effect being due to the sample size.

Overall these studies support that oral NR is available to human skeletal muscle, and reveal novel anti-inflammatory NR properties, both of which may be beneficial in the context of aging, muscle, or inflammatory disease groups.

## Author Contributions

Y.S.E and G.G.L conceived the trial. Y.S.E, K.N.M, A.P and G.G.L designed the trial. Y.S.E conducted the clinical trial. Y.S.E, K.K, D.A.T, and A.G conducted and supported the high-resolution mitochondrial respirometry. I.A performed bioinformatic and molecular pathway analyses. C.L.D, D.M.C, L.O, AS, and YL contributed to molecular analysis. G.W contributed helpful discussions and supported blood analyses. R.S.F, M.S.S, and C.B. undertook targeted metabolomics and quantitation of NAD^+^-related metabolites, while Y.S.E analyzed the data. C.V.B, N.J, M.W, and S.J.E.L conducted the ^31^P MRS *in vivo* NAD^+^ studies and analyzed the data. Y.S.E and P.N conducted statistical analyses. Y.S.E, C.B, and G.G.L took the leading role in writing the manuscript and creating the figures. All authors contributed to the editing and proofreading of the final draft.

## Acknowledgements

This work was supported by an MRC PhD studentship (YSE), Medical Research Council (MRC) Confidence in Concept (CiC) award (AP and GGL, CiC4/21), Wellcome Trust Senior Fellowship (GGL, 104612/Z/14/Z), Marie Sklodowska-Curie grant (AG, No 705869), and the Roy J. Carver Trust and National Institutes of Health (CB, R01HL147545). We thank ChromaDex (Irvine, California) for providing nicotinamide riboside and placebo capsules. We also thank all staff at the NIHR/Wellcome Trust Clinical Research Facility - Queen Elizabeth Hospital Birmingham. We acknowledge Professor Alexandra Sinclair, Professor Jeremy Tomlinson, Dr Zaki Hassan-Smith, and Dr Keira Markey for their helpful discussions, Dr Alpesh Thakker for his input into high-resolution respirometry, and Professor Janet Lord for her help with recruitment of participants.

## METHODS

### Study conduct

**NR supplementation study:** the study conducted between July 2016 and August 2017 at the National Institute for Health Research/Wellcome Trust Clinical Research Facility at the Queen Elizabeth Hospital Birmingham, UK. The Solihull NRES Committee gave ethical approval (REC reference number 16/WM/0159). All participants provided written informed consent. The study was registered on www.clinicaltrials.gov (Identifier: NCT02950441).

**^31^P MRS study:** the study was conducted between November 2017 and February 2018 at the Birmingham University Imaging Centre, University of Birmingham. The study was approved by the University of Birmingham Research Ethics Committee (ERN_11-0429AP67). All participants provided written informed consent.

The studies were undertaken according to the principles of the Declaration of Helsinki and followed the Guidelines for Good Clinical Practice.

### Study population

**NR supplementation study:** aged volunteers were recruited from the Birmingham 1000 Elders group (https://www.birmingham.ac.uk/research/activity/mds/centres/healthy-ageing/elders.aspx). All participants fulfilled the inclusion criteria including: male sex, age 70– 80 years, BMI 20 – 30 kg/m^2^, able to discontinue aspirin for 3 days prior to the muscle biopsy, and able to discontinue statins and vitamin D supplements for a week before the study and for the duration of the study. Exclusion criteria included: serious active medical conditions including inflammatory diseases or malignancies, significant past medical history including diabetes mellitus, ischaemic heart disease, cerebrovascular disease, respiratory disease requiring medication, or epilepsy, blood pressure >160/100mmHg, or treatment with oral anti-coagulants.

**^31^P MRS study:** young (20 - 35 years; n=16) and aged (60 – 75 years; n = 11) subjects were recruited through public adverts and a recruitment database held by the School of Psychology, University of Birmingham. Inclusion criteria in both age groups were: BMI 20 - 30 kg/m^2^, and absence of a history of metabolic, cardiovascular or respiratory disease.

### Study design

**NR supplementation study:** single centre, double blind, placebo-controlled, and crossover study. Aim was to obtain complete assessments from 12 aged individuals. Participants attended for a screening visit (visit 1) when an informed written consent was obtained after ensuring they fulfil all inclusion criteria. For all subsequent study visits (2 to 5), the participants attended at 08:00 in a fasting state from the night before. Regarding the post-interventions visits (3 and 5), the participants took the last NR/placebo dose 14 hours prior to the assessments.

**^31^P MRS study:** participants were requested to avoid exercise and alcohol 24 hours prior to the visit, and avoid caffeine 12 hours prior to the visit. Due to ethical considerations, participants were not required to undergo both the muscle and brain assessments. The aim was to recruit a minimum of 10 participants in each group.

### Randomization and blinding

**NR supplementation study:** Participants were allocated to either NR or placebo. A randomization list was held by the clinical trials pharmacist at the clinical research facility. The study investigators, nurses, and participants were all blinded to the intervention allocation during the trial.

### Intervention

**NR supplementation study:** NR was supplied as 250 mg capsules by the manufacturer (Niagen®, ChromaDex, Irvine, CA). Participants received NR 500mg twice daily or matched placebo for 21 days with 21 days washout period between the NR and placebo periods (Suppl. Figure 1A).

### Assessments

**NR supplementation study:** assessments undertaken in each study visits are detailed in Suppl. Figure 1.

**^31^P MRS study:** each of the muscle and brain measurements were undertaken in one day.

## METHOD DETAILS

### Blood pressure

Blood pressure (Welch Allyn, USA) was measured at the start of each visit after an overnight fast. At the trial visits, participants rested for 15mins in a supine position before blood pressure was measured. An appropriately sized cuff was selected to encircle at least 80% of the arm and the same was used every visit for all participants, and on the same arm. Blood pressure was measured in triplicates and the mean was recorded.

### Hand-held dynamometry

Peak absolute strength (kilograms) and relative handgrip strength (kilograms of force per kilogram of body weight) were measured in triplicate bilaterally using a dynamometer (Takei Instruments, Japan). The highest measurement values were included for analysis.

### Muscle biopsies

Resting vastus lateralis muscle biopsies were obtained from 12 men by a single investigator (Y.S.E) using a percutaneous Bergstrom technique as previously described (Bergstrom, 1975) under local anaesthesia (1% lignocaine). The biopsy sample (100 – 150mg) was immediately dried on clean filter paper and approximately 10mg of tissue was cut and placed on ice cold BIOPS buffer for high resolution respirometry (see below). The rest of the muscle tissue was immediately snap frozen in liquid nitrogen and stored at −80°C pending analysis.

### Indirect calorimetry

Participants were allowed to rest for 60 mins after insertion of the arterial and venous catheters. Then they lay supine in a comfortable position while a transparent ventilated canopy was placed over their head. Plastic sheet attached to the hood was placed around the subjects to form a seal. The room temperature, barometric pressure, and humidity were measured by a hygrometer (Oregon Scientific). During the measurement period, participants remained supine, and breathed normally. Respiratory measurements, including resting oxygen consumption (VO_2_) and carbon dioxide production (VCO_2_) and respiratory exchange ratio (VCO_2_/VO_2_), using the mixing chamber mode of the metabolic cart (AEI MOXUS II Metabolic System). Measurements were collected at fasting and then every 30 minutes for 2 hours following a 75g oral glucose load. Measurements for each period lasted 15 mins so the first and last minutes could be discarded, and the mean value for the middle 5 minutes was recorded.

### Arterio-venous difference technique

An arterial catheter was inserted into a radial artery and a retrograde cannula was inserted into in a deep antecubital vein draining a forearm muscle, on the opposite side of the arterial line. To prevent contamination of the muscle venous blood with the mixed blood from the hand, a wrist cuff was inflated to 200mmHg for 3 mins before sampling. Blood sampling was undertaken simultaneously from the arterial and venous sites at fasting and every 20mins following oral glucose load for 120 min.

### Venous occlusive plethysmography

Forearm muscle blood flow was measured by venous occlusive plethysmography (Hokanson, USA) (Wythe *et al*., 2015) as previously described (Greenfield, Whitney and Mowbray, 1963). Blood flow measurements were taken immediately after each blood sampling.

### NAD^+^ metabolomics

Muscle tissue was pulverised and approximately 10 mg was used for each of the acid (A) and basic (B) metabolite extraction. For each sample, internal standard mixtures for each of A and B were prepared. The extraction was undertaken using 0.2 ml of ice-cold LC-MS/MS grade methanol and kept on ice before adding 0.3ml of internal standard made in LC-MS grade water. Samples were sonicated in an acetone water bath (at −4 **°**C) for 20 seconds, placed back on ice, and then incubated at 85 **°**C with constant shaking at 1050 rpm for 5 minutes. Samples were then placed on ice for 5 min and centrifuged (16.1k x g, 10 minutes, 4 °C). The supernatant was transferred to clean tubes and dried using a speed vacuum. The dried extract was re-suspended in 30 µl of either LC-MS grade water for acid extract or 10 mM ammonium acetate for alkaline extract and centrifuged (16.1k x g, 3 minutes, 4 °C). The supernatant was carefully transferred to a Waters Polypropylene 0.3 ml plastic screw-top vial. The pellet was then dried using a speed vacuum, pellet was weighed, and later used to normalize data that were finally reported as pmol/mg.

Otherwise, muscle, blood and urine metabolomics were undertaken as previously described (Trammell and Brenner, 2013; S. A. Trammell *et al*., 2016).

### Blood biochemical analyses

Blood was drawn from the arterial and venous catheters into heparinised blood tubes. Plasma was rapidly separated by centrifugation at 4°C and was then snap frozen. Plasma glucose, NEFA, and lactate concentrations were measured using commercially available kits on an ILAB 650 Clinical Chemistry Analyser (Werfen Ltd, UK). Insulin was measured using commercially available assay as per the manufacturer’s instructions (Mercodia, Sweden). Homeostasis model assessment of insulin resistance (HOMA-IR) was calculated using the formula [fasting glucose (mmol/L) × fasting insulin (mU/L)/22.5].

Lipid profile, urea and electrolytes, and thyroid function tests were all measured on the Roche Modular Platforms (Roche, Switzerland). Full blood count was measured on a Beckman Coulter DxH analyser (USA).

### High resolution respirometry on permeabilized muscle fibres

*Ex vivo* mitochondrial function was determined by measuring oxygen consumption polarographically using a two-chamber Oxygraph-2k (OROBOROS Instruments). Oxygen consumption reflects the first derivative of the oxygen concentration (nmol/ml) in time in the respiration chambers and is termed oxygen flux [pmol/(s*mg)], corrected for wet weight muscle tissue (2–5 mg) introduced into the chamber. Measurements were undertaken according to a previously described protocol (Pesta and Gnaiger, 2012). Similar results were obtained if respiration rates were corrected for mitochondrial DNA (mtDNA) copy number or citrate synthase activity.

### Mitochondrial density assessments

For citrate synthase activity, 5mg of snap frozen human muscle was used and the measurement was undertaken as previously described (Horscroft *et al*., 2015). Mitochondrial DNA (mtDNA) copy number was determined using quantitative real time PCR. mtDNA copy number was calculated from the ratio of NADH dehydrogenase subunit 1 (ND1) to lipoprotein lipase (LPL) (mtDNA/nuclear DNA) as previously described (Phielix *et al*., 2008).

### RNA sequencing

RNA was extracted from frozen muscle tissue using Tri Reagent (Sigma-Aldrich) following manufacturer’s instructions. RNA-sequencing was performed using libraries prepared with Lexogen Quantseq FWD library preparation kit and essentially analysed as previously described (Akerman *et al*., 2017). A detailed description is provided in the supplemental methods section.

### Protein immunoblotting

Muscle biopsies were homogenised in ice-cold sucrose lysis buffer (50 mM Tris/HCl (pH 7.5), 250 mM Sucrose, 10mM Na-β-Glycerophosphate, 5mM Na-Pyrophosphate, 1mM Benazmidine, 1 mM EDTA, 1 mM EGTA, 1% Triton X-100, 1 mM Na3VO4, 50 mM NaF, 0.1% β-Mercaptoethanol, supplemented with protease inhibitor cocktail). Samples (40-100µg of protein extract) were loaded into 4-15% Tris/Glycine precast gels (BioRad) prior to electrophoresis. Proteins were transferred onto PVDF membranes (Millipore) for 1h at 100V. A 5% skimmed milk solution made up with Tris-buffered saline Tween-20 (TBS-T, 0.137M NaCl, 0.02M Tris-base 7.5pH, 0.1% Tween-20) was used to block each membrane for 1h before being incubated overnight at 4°C with appropriate primary antibodies. Membranes were washed in TBS-T three times prior to incubation in horse radish peroxidase-conjugated secondary antibody at room temperature for 1h. Membranes were then washed in TBS-T prior to antibody detection via enhanced chemiluminescence horseradish peroxidase substrate detection kit (Millipore). Images were undertaken using a G:Box Chemi-XR5 (Syngene).

### Inflammatory cytokines

We performed a multiplex cytokine bead assay using the Bio-Plex Pro Human Cytkine 17-plex panel analyzed with a flow-cytometry based Luminex 200 reader. The levels of IL-1b, IL-2, IL-4, IL-5, IL-6, IL-7, IL-8, IL-10, IL-12, IL-13, IL-17, G-CSF, GM-CSF, IFN-g, MCP-1, MIP-1b, and TNF-a were measured on the participants’ sera as per the manufacturer’s instructions. Only IL-2, IL-5, IL-6, IL-8, IL-12, IFN-g, TNF-a, MCP-1, and MIP-1b were within detection range. High sensitive CRP was measured using CRPHS: ACN 217 on COBAS 6000 analyser (Roche, USA). All measurements were undertaken in duplicates.

### In vivo ^31^P MRS measurements of skeletal muscle and brain

In-vivo ^31^P MRS was performed on a 3T Philips Achieva MRI system with the Philips P-140 surface coil. Localisation was based on the sensitivity profile of the coil and spectral data was acquired over 2048 complex points at a sampling frequency of 3000Hz with a repetition time of 3500ms. ^1^H broadband decoupling was applied during acquisition. For calf measurements 256 averages were collected. For brain measurements, the coil was positioned in the region of occipital lobe and 768 averages were collected. An in-house pre-processing and spectral fitting pipeline was used for analysis. Metabolite simulation and least-squares fitting steps were implemented as described previously (Lu *et al*., 2014) and metabolite concentrations were calculated by normalising to an assumed α-ATP concentration of 2.8mM.

Measurement repeatability was tested by scanning the same subject three times with identical positioning and sequence parameters. Subsequently the subject was removed and repositioned and a fourth measurement made. Coefficients of variation of 3% and 9% were found across the four repeat measures of NADH+NAD^+^ for brain and calf, respectively.

## QUANTIFICATION AND STATISTICAL ANALYSIS

Sample size for this experimental medicine study was decided upon based on previous experience from studies using the same methodological design, whereby the proposed sample size was sufficient to detect significant differences at the 5% level. The analysis was based on data from all participants who were randomized, and completed all the study visits and assessments. Outcome data were reported as mean ± SEM (or median and quartiles where appropriate). In the NR supplementation study, comparisons of participants between placebo and NR supplementation phases were undertaken using paired t-tests. In addition, further data analysis taking into account the period effect was undertaken, by grouping the participants into those who were randomised to NR first and second. This is to look for carryover effect across all analyses. Wherever relevant, area under the curve was calculated using the trapezoid method. For the ^31^P MRS study, young and aged groups were compared using an independent sample t-test. Data were analyzed using IBM SPSS Statistics version 22 and GraphPad Prism version 7.0.

## SUPPLEMENTAL INFORMATION

**Suppl. Table 1.** Cardiometabolic parameters at baseline and after nicotinamide riboside (NR) and placebo. Data are presented as median (1^st^ quartile, 3^rd^ quartile)

| Parameter | Baseline | NR | Placebo |
| --- | --- | --- | --- |
| Age (years) | 75 (72, 78) | - | - |
| Weight (kg) | 82.6 (76.5, 90.1) | 83 (76.3, 89.9) | 82.6 (77.0, 89.6) |
| BMI (kg/m<sup>2</sup>) | 26.6 (25.0, 30.0) | 26.9 (24.9, 29.5) | 26.9 (25.1, 29.3) |
| Systolic blood pressure (mmHg) | 139 (136, 154) | 138 (130, 147) | 134 (126, 155) |
| Diastolic blood pressure (mmHg) | 86 (83, 93) | 84 (79, 93) | 81 (77, 90) |
| Fasting glucose (mmol/L) | 5.88 (5.63, 6.31) | 6.04 (5.81, 6.23) | 5.85 (5.67, 6.30) |
| Fasting insulin (mU/L) | 7.07 (7.27, 8.67) | 6.27 (5.47, 7.07) | 7.06 (5.47, 8.67) |
| Cholesterol (mmol/L) | 4.8 (3.6, 5.2) | 4.6 (3.9, 5.6) | 4.6 (4.2, 5.3) |
| HDL cholesterol (mmol/L) | 1.4 (1.2, 1.5) | 1.3 (1.2, 1.5) | 1.4 (1.1, 1.5) |
| Triglycerides (mmol/L) | 1.0 (0.6, 1.2) | 1.0 (0.8, 1.2) | 1.0 (0.7, 1.2) |

**Suppl. Table 2.**
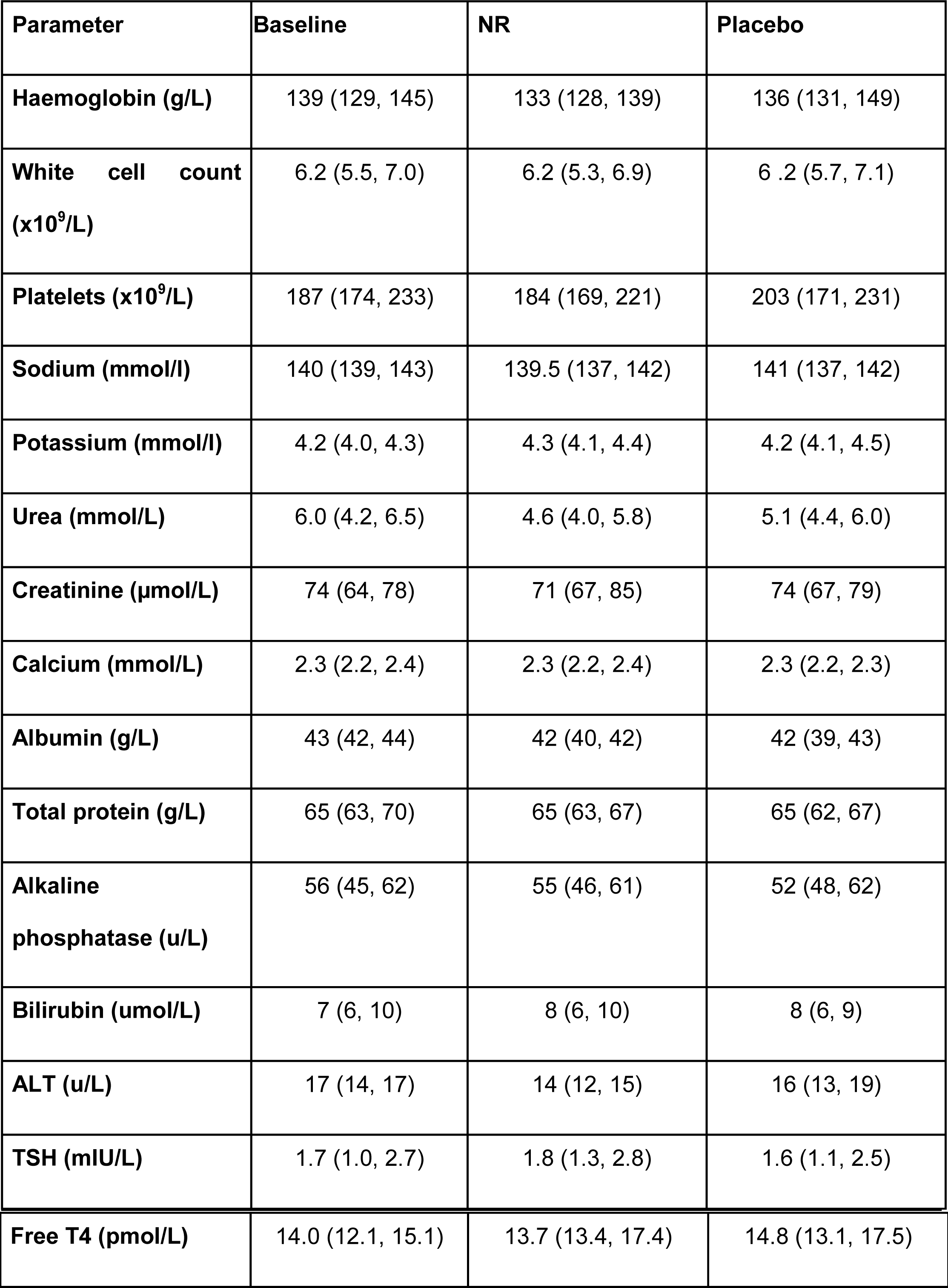
Safety parameters at baseline and after nicotinamide riboside (NR) and placebo. Data presented as median (1^st^ quartile, 3^rd^ quartile). ALT, alanine aminotransferase; TSH, thyroid stimulating hormone; Free T4, free thyroxine

**Suppl. Table 3 (EXCEL file).** NAD+ metabolomics data in skeletal muscle, blood and urine after each of the baseline, nicotinamide riboside (NR), placebo, and washout phases.

**Suppl. Table 4 (EXCEL file).** List of genes up- and downregulated in skeletal muscle biopsies upon nicotinamide riboside supplementation (N = 12).

**Suppl. Table 5 (EXCEL file).** List of genes in Gene Ontology terms related to processes up- and downregulated in skeletal muscle biopsies upon nicotinamide riboside supplementation (N = 12).

**Suppl. Figure 1.**
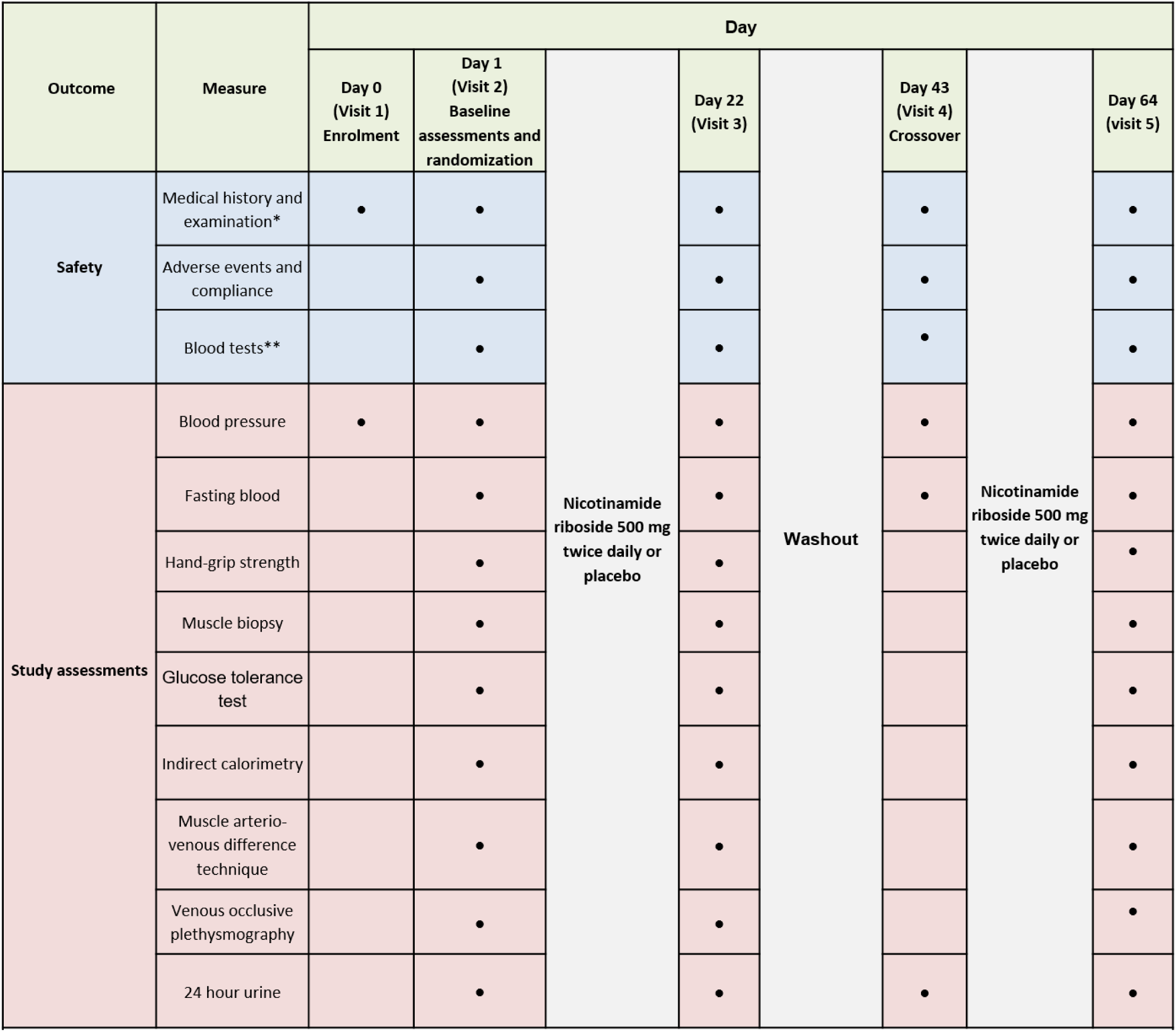
Outline of the measures undertaken in each of the 5 study visits and the time interval between visits. * Body weight and height, systemic examination, and resting electrocardiogram. ** Full blood count, and renal, liver, and thyroid functions.

**Suppl. Figure 2.**
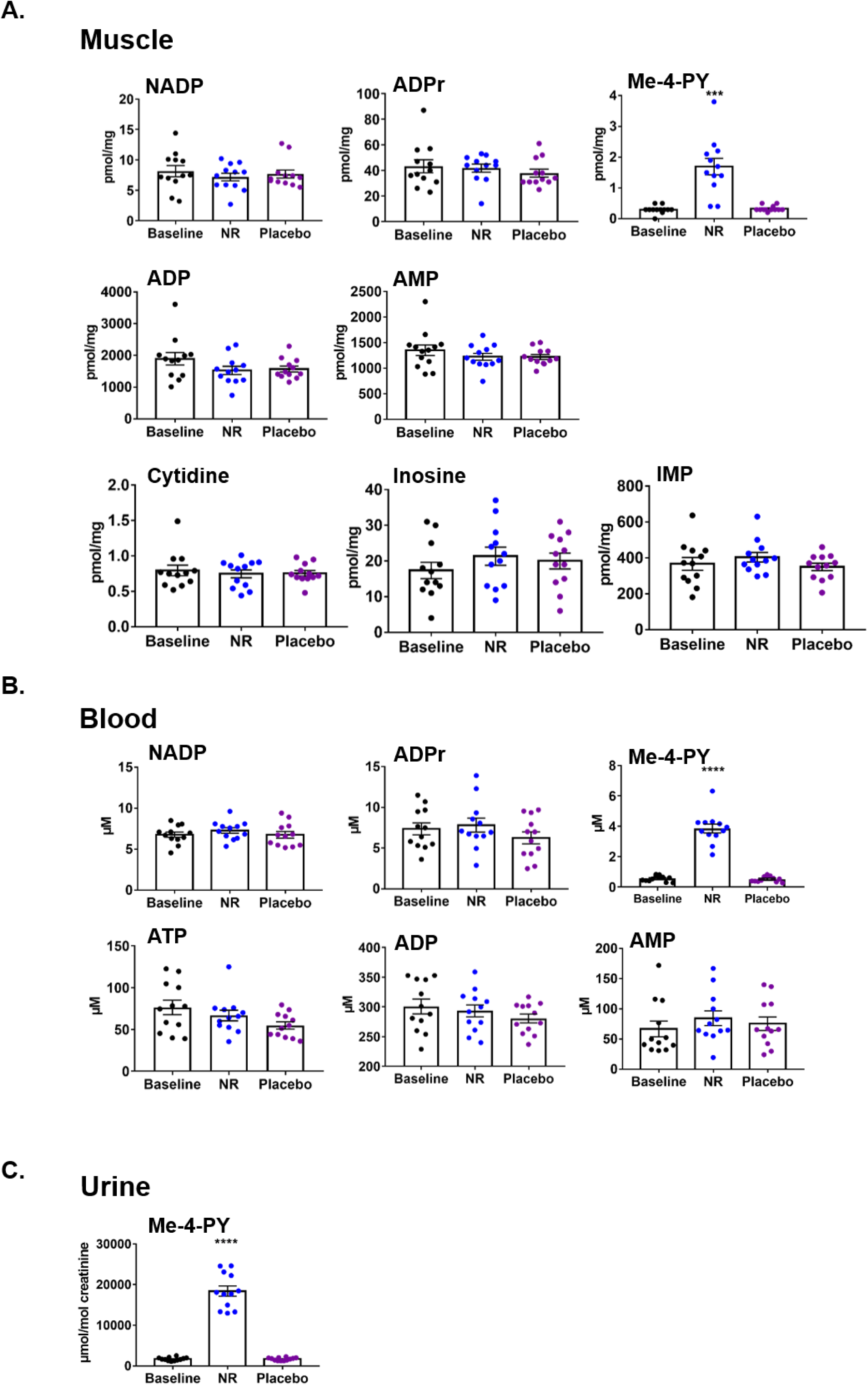
Remainder of LC-MS/MS NAD^+^ metabolomics in (A) skeletal muscle, (B) whole blood, and (C) urine, which were not shown in Figure 1. NADP, nicotinamide adenine dinucleotide phosphate; ADPr, adenosine diphosphate ribose, Me-4-py, N1-Methyl-4-pyridone-5-carboxamide; ATP, adenosine triphosphate; ADP, adenosine diphosphate; AMP, adenosine monophosphate. Data are obtained from 12 participants at each phase and presented as mean ± SEM. Significance was set at p < 0.05 using paired t-test. The absence of significance symbols indicates lack of statistical significance.

**Suppl. Figure 3.**
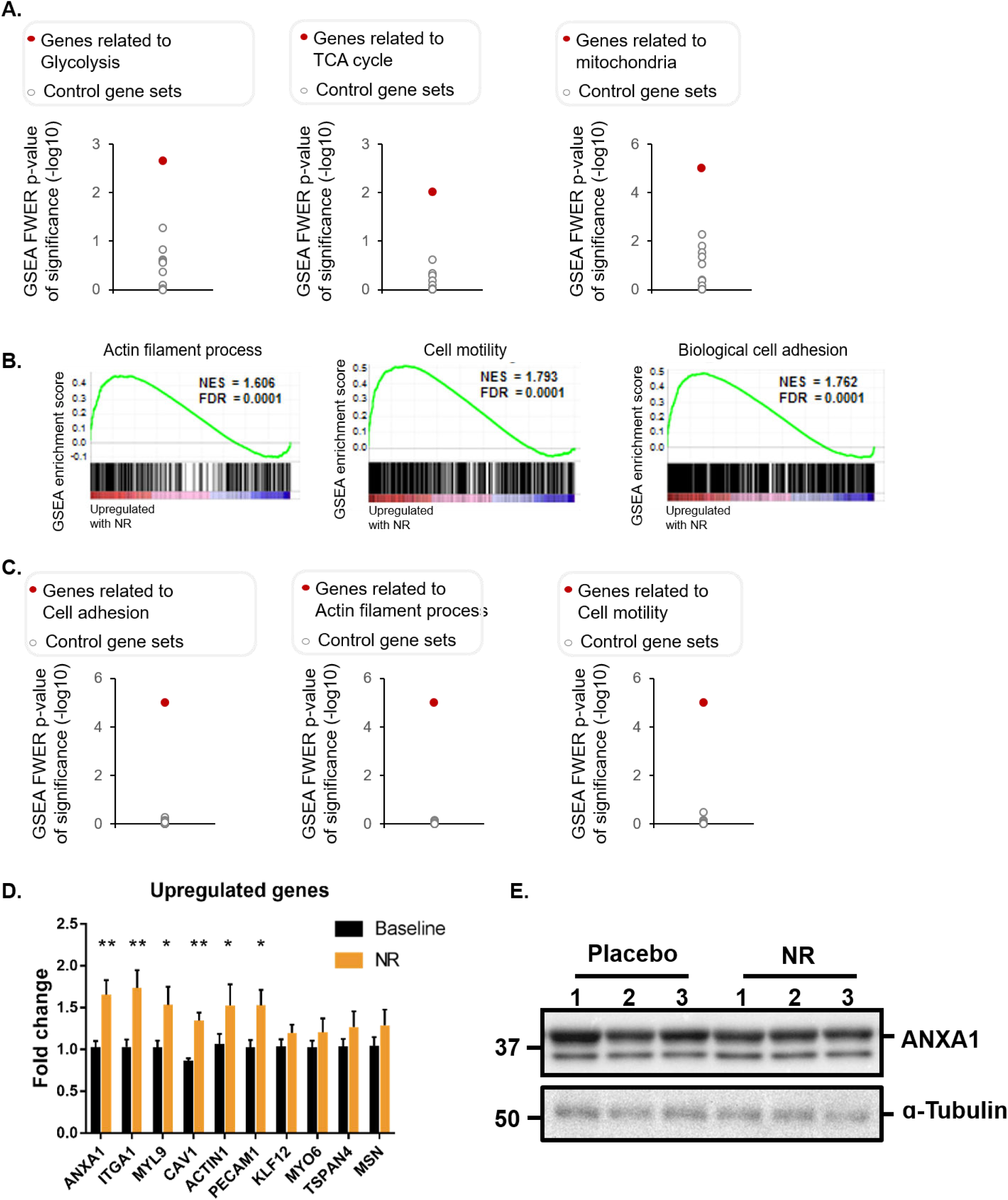
(A) Bar plots show GSEA p-value of significance (-Log10) for enrichment of genes belonging to the pathways gylcolysis, TCA cycle and mitochondria for downregulated targets of NR supplementation (in red). The same analysis on 10 gene sets of equal size and expression level do not reveal enrichment amongst downregulated targets of NR (in grey) (B) Gene set enrichment analysis (GSEA) suggests that genes belonging to the gene sets “Actin filament process”, “Cell motility”, and “Biological cell adhesion” are upregulated upon NR supplementation. Normalized enrichment score (NES) and nominal p-value is presented on the top left corner of the graph. (C) Bar plots show GSEA p-value of significance (-Log10) for enrichment of genes belonging to the pathways “Actin filament process”, “Cell motility”, and “Biological cell adhesion” for upregulated targets of NR supplementation (in red). The same analysis on 10 gene sets of equal size and expression level do not reveal enrichment amongst upregulated targets of NR (ingrey). (D) Quantitative PCR analysis of a select panel of upregulated genes identified through differential gene expression analysis. GAPDH was used as housekeeping gene. Error bars represent SEM (n=12). (E) Quantification of Annexin A1 (ANXA1) protein using immunoblotting assay. Tubulin was used as a loading control.

**Suppl. Figure 4.**
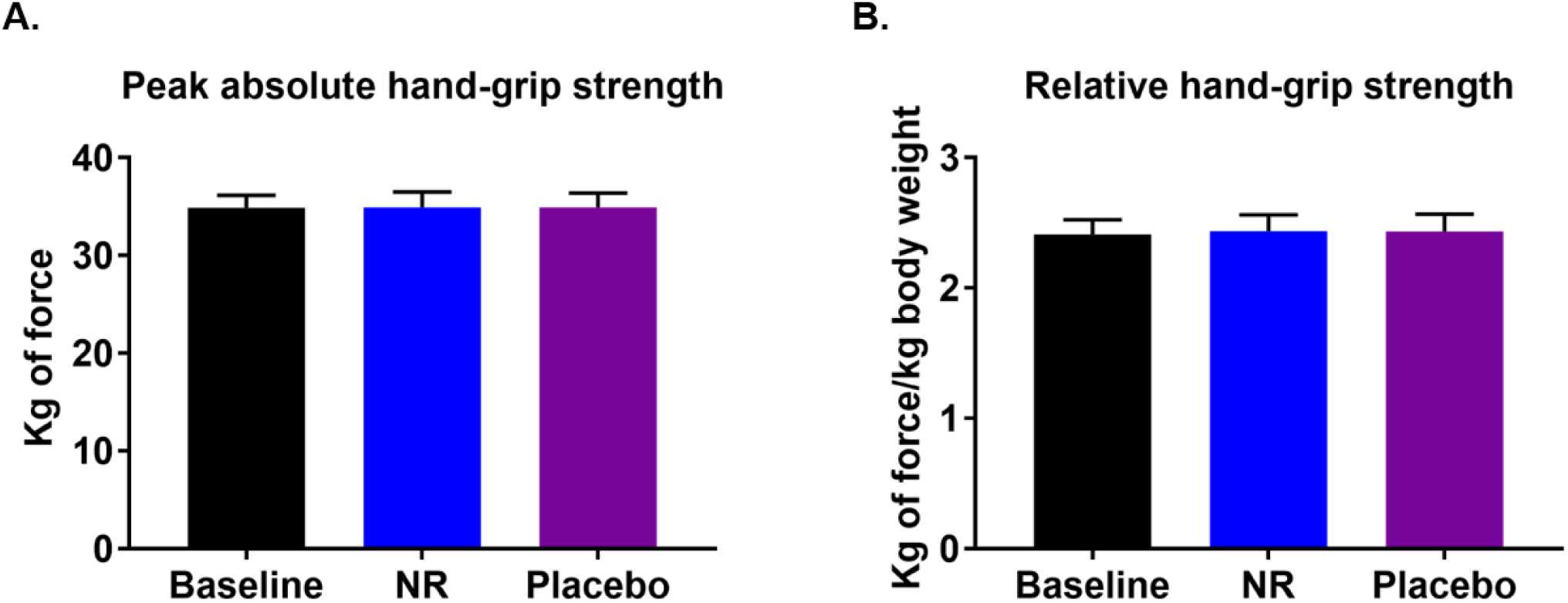
(A) Peak hand-grip strength at baseline and after each of the nicotinamide riboside (NR) and placebo phases. (B) Similar to (A) but data presented relative to body weight. Data are obtained from 12 participants at each phase and presented as mean ± SEM. Significance was set at p < 0.05 using paired t-test. The absence of significance symbols indicates lack of statistical significance.

**Suppl. Figure 5.**
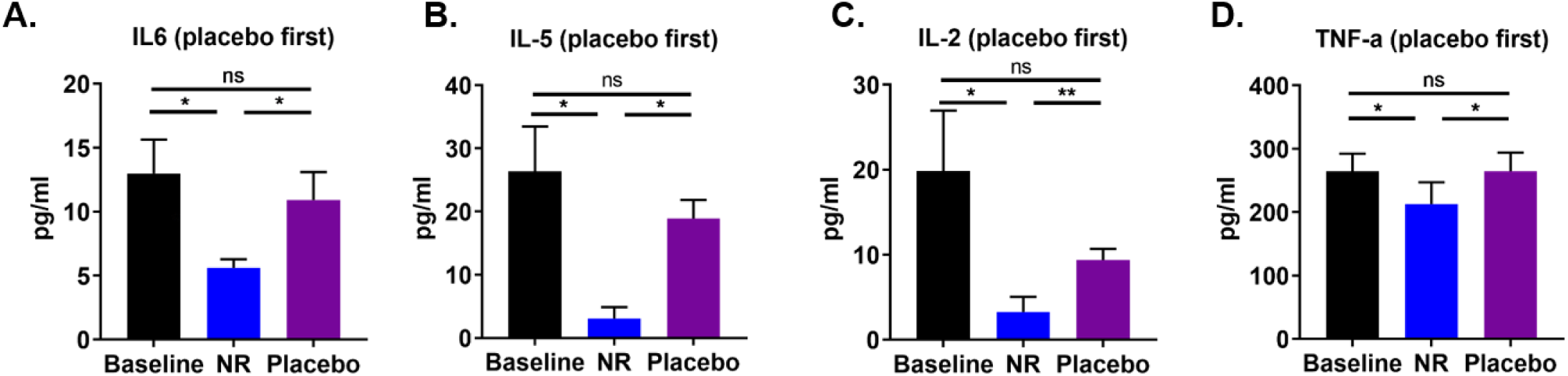
Levels of serum inflammatory cytokines at baseline and after each of the nicotinamide riboside (NR) and placebo phases, including: (A) interleukin 6 (IL-6), (B) interleukin 5 (IL-5), (C) interleukin 2 (IL-2), and (D) tumor necrosis factor-alpha (TNF-a). Here period effect analysis is shown and each panel is produced from only the 6 subjects that were randomized to placebo first as a demonstration of the NR carryover effect, evident in the cases IL-2 (C) and TNF-a (D). Data are presented as mean ± SEM. Significance was set at p < 0.05 using paired t-test.

## SUPPLEMENTAL METHODS

### RNA sequencing

RNA was extracted from frozen muscle tissue using Tri Reagent (Sigma-Aldrich) following manufacturer’s instructions. Sequencing libraries were prepared using RNA (RIN >7) with the Lexogen Quantseq3 FWD kit. Libraries were sequenced using HiSeq2000 across 4 flowcells generating 75bp long single ended reads (average read depth of 6-10M/sample, which is higher than the 4M reads / sample required for analysis for this type of library). All samples were prepared and sequenced as a single pool. Trimmomatic software (v0.32) and bbduk.sh script (Bbmap suite) were used to trim the ILLUMINA adapters, polyA tails and low quality bases from reads. Trimmed reads were then uniquely aligned to the human genome (hg38) using STAR with default settings (v2.5.2b) and the Gencode (v28, Ensembl release 92) annotation as the reference for splice junctions. Mapped reads were quantified using HT-seq (v0.9.1) using Gencode (v28) genes (-intersection-nonempty flag). Differential gene expression was obtained using DEseq2 with paired baseline and treatment samples. In this analysis we did not use a cutoff to remove lowly expressed genes. Inclusion of lowly expressed genes (at arbitrary cut-offs) had little bearing on our results (97.8% of differentially expressed genes at p < 0.05 were identical between no cutoff, and a cut-off of > 3). Of note, volcano plot was drawn with a cut-off (>3) in order to visualise the typical “V” shape using R. Differentially expressed genes between baseline (control) and NR treated samples at p-value = <0.05 were annotated using Biological processes (BP) gene sets with DAVID tool (Huang, Brad T Sherman, and Lempicki 2009; Huang, Brad T. Sherman, and Lempicki 2009). We obtained similar results using gene annotation tool within Gene Set Enrichment Analysis (GSEA) suite (Liberzon et al. 2015; Subramanian et al. 2005) for gene sets from KEGG pathways and C5-Biological processes. In addition, we have used GSEA analysis tool to interrogate specific gene sets against our pre-ranked expression data (Control vs NR treatment). GSEA calculates an Enrichment

Score (ES) by scanning a ranked-ordered list of genes (according to significance of differential expression (-log10 p-value), increasing a running-sum statistic when a gene is in the gene set and decreasing it when it is not. The top of this list (red) contains genes upregulated upon NR+ treatment while the bottom of the list (blue) represents downregulated genes. Each time a gene from the interrogated gene set (i.e. Glycolysis, mitochondria, TCA cycle) is found along the list, a vertical black bar is plotted (hit). If the “hits” accumulate at the bottom of the list, then this gene set is enriched in upregulated genes (and vice versa). If interrogated genes are distributed homogenously across the rank ordered list of genes, then that gene set is not enriched in any of the gene expression profiles (i.e. control gene sets of similar expression levels to interrogated gene sets). GSEA was used in pre-ranked mode with parameters -norm meandiv -nperm 1000 - scoring_scheme weighted. 10 gene sets of equal size and similar expression levels to the interrogated gene sets were generated using a custom pipeline in R (available upon request). We have interrogated the following gene sets: GO0048870; cell motility, GO0030029; actin filament based process, GO0022610; Biological cell adhesion, (also GO0007155 cell adhesion with similar results), M15112: Wong Mitochondria gene module, M3985: KEGG citrate cycle TCA cycle, merge of M15109: BIOCARTA Glycolysis pathway and M5113: REACTOME glycolysis.

## References

1. Airhart, S. E. et al. (2017) ‘An open-label, non-randomized study of the pharmacokinetics of the nutritional supplement nicotinamide riboside (NR) and its effects on blood NAD+ levels in healthy volunteers’, PLOS ONE, 12(12), p. e0186459.

2. Akerman, I. et al. (2017) ‘Human Pancreatic β Cell lncRNAs Control Cell-Specific Regulatory Networks’, Cell Metabolism, 25(2), pp. 400–411.

3. Amici, S. A. et al. (2018) ‘CD38 Is Robustly Induced in Human Macrophages and Monocytes in Inflammatory Conditions’, Frontiers in Immunology, 9, p. 1593.

4. Bergstrom, J. (1975) ‘Percutaneous Needle Biopsy of Skeletal Muscle in Physiological and Clinical Research’, Scandinavian Journal of Clinical & Laboratory Investigation, 35(7), pp. 609–616.

5. Bickerton, A. S. T. et al. (2007) ‘Preferential uptake of dietary fatty acids in adipose tissue and muscle in the postprandial period’, Diabetes, 56(1), pp. 168–176.

6. Bieganowski, P. and Brenner, C. (2004) ‘Discoveries of nicotinamide riboside as a nutrient and conserved NRK genes establish a preiss-handler independent route to NAD+ in fungi and humans’, Cell, 117(4), pp. 495–502. doi: 10.1016/S0092-8674(04)00416-7.

7. Bogan, K. L. and Brenner, C. (2008) ‘Nicotinic acid, nicotinamide, and nicotinamide riboside: a molecular evaluation of NAD+ precursor vitamins in human nutrition.’, Annual review of nutrition, 28, pp. 115–130.

8. Brown, K. D. et al. (2014) ‘Activation of SIRT3 by the NAD+ precursor nicotinamide riboside protects from noise-induced hearing loss’, Cell Metabolism, 20(6), pp. 1059–1068.

9. Camacho-Pereira, J. et al. (2016) ‘CD38 Dictates Age-Related NAD Decline and Mitochondrial Dysfunction through an SIRT3-Dependent Mechanism’, Cell Metabolism, 23(6), pp. 1127–1139.

10. Cantó, C. et al. (2012) ‘The NAD+ precursor nicotinamide riboside enhances oxidative metabolism and protects against high-fat diet-induced obesity’, Cell Metabolism, 15(6), pp. 838–847.

11. Chaleckis, R. et al. (2016) ‘Individual variability in human blood metabolites identifies age-related differences’, Proceedings of the National Academy of Sciences, 113(16), pp. 4252– 4259.

12. Clement, J. et al. (2018) ‘The Plasma NAD+ Metabolome is Dysregulated in “normal” Ageing’, Rejuvenation Research, p. rej.2018.2077.

13. Costford, S. R. et al. (2010) ‘Skeletal muscle NAMPT is induced by exercise in humans’, AJP: Endocrinology and Metabolism, 298(1), pp. E117–E126.

14. Covarrubias, A. J. et al. (2019) ‘Aging-related inflammation driven by cellular senescence enhances NAD consumption via activation of CD38 + macrophages’, bioRxiv. doi: 10.1101/609438.

15. Cruz-Jentoft, A. J. et al. (2010) ‘Sarcopenia: European consensus on definition and diagnosis: Report of the European Working Group on Sarcopenia in Older People’, Age and Ageing, 39(4), pp. 412–423.

16. Das, A. et al. (2018) ‘Impairment of an Endothelial NAD+-H2S Signaling Network Is a Reversible Cause of Vascular Aging.’, Cell, 173(1), p. 74–89.e20.

17. Diguet, N. et al. (2018) ‘Nicotinamide Riboside Preserves Cardiac Function in a Mouse Model of Dilated Cardiomyopathy’, Circulation, 137(21), pp. 2256–2273.

18. Dodds, R. M. et al. (2014) ‘Grip Strength across the Life Course: Normative Data from Twelve British Studies’, PLoS ONE, 9(12), p. e113637.

19. Dollerup, O. L. et al. (2018) ‘A randomized placebo-controlled clinical trial of nicotinamide riboside in obese men: safety, insulin-sensitivity, and lipid-mobilizing effects’, The American Journal of Clinical Nutrition, 108(2), pp. 343–353.

20. Elhassan, Y. S., Philp, A. A. and Lavery, G. G. (2017) ‘Targeting NAD+ in Metabolic Disease: New Insights Into an Old Molecule’, Journal of the Endocrine Society, 1(7), pp. 816–835.

21. Fang, E. F. et al. (2017) ‘NAD + in Aging: Molecular Mechanisms and Translational Implications’, Trends in Molecular Medicine, 23(10), pp. 899–916.

22. Fletcher, R. S. et al. (2017) ‘Nicotinamide riboside kinases display redundancy in mediating nicotinamide mononucleotide and nicotinamide riboside metabolism in skeletal muscle cells’, Molecular Metabolism, 6(8), pp. 819–832.

23. Frederick, D. W. et al. (2016) ‘Loss of NAD Homeostasis Leads to Progressive and Reversible Degeneration of Skeletal Muscle’, Cell Metabolism, 24(2), pp. 269–282.

24. Gariani, K. et al. (2016) ‘Eliciting the mitochondrial unfolded protein response by nicotinamide adenine dinucleotide repletion reverses fatty liver disease in mice’, Hepatology, 63(4), pp. 1190–1204. doi: 10.1002/hep.28245.

25. Gomes, A. P. et al. (2013) ‘Declining NAD+ induces a pseudohypoxic state disrupting nuclear-mitochondrial communication during aging’, Cell, 155(7), pp. 1624–1638.

26. Goody, M. F. et al. (2010) ‘Nrk2b-mediated NAD+ production regulates cell adhesion and is required for muscle morphogenesis in vivo. Nrk2b and NAD+ in muscle morphogenesis’, Developmental Biology, 344(2), pp. 809–826.

27. Goody, M. F. and Henry, C. A. (2018) ‘A need for NAD+ in muscle development, homeostasis, and aging’, Skeletal Muscle, 8(1), pp. 1–14.

28. Greenfield, A. D. M., Whitney, R. J. and Mowbray, J. F. (1963) ‘Methods for the investigation of peripheral blood flow’, Brit Med Bull, 19(2), pp. 101–109.

29. Hagopian, K., Ramsey, J. J. and Weindruch, R. (2003) ‘Influence of age and caloric restriction on liver glycolytic enzyme activities and metabolite concentrations in mice’, Experimental Gerontology, 38(3), pp. 253–266.

30. Horscroft, J. A. et al. (2015) ‘Altered Oxygen Utilisation in Rat Left Ventricle and Soleus after 14 Days, but Not 2 Days, of Environmental Hypoxia’, PLOS ONE, 10(9), p. e0138564.

31. Huang, D. W., Sherman, B. T. and Lempicki, R. A. (2009) ‘Bioinformatics enrichment tools: paths toward the comprehensive functional analysis of large gene lists’, Nucleic Acids Research, 37(1), pp. 1–13.

32. Huang, D. W., Sherman, B. T. and Lempicki, R. A. (2009) ‘Systematic and integrative analysis of large gene lists using DAVID bioinformatics resources’, Nature Protocols, 4(1), pp. 44–57.

33. Imai, S. and Yoshino, J. (2013) ‘The importance of NAMPT/NAD/SIRT1 in the systemic regulation of metabolism and ageing’, *Diabetes*, Obesity and Metabolism, 15(S3), pp. 26–33.

34. Ingram, D. K. and Roth, G. S. (2011) ‘Glycolytic inhibition as a strategy for developing calorie restriction mimetics’, Experimental Gerontology. Elsevier Inc., 46(2–3), pp. 148–154.

35. Kannt, A. et al. (2015) ‘Association of nicotinamide-N-methyltransferase mRNA expression in human adipose tissue and the plasma concentration of its product, 1-methylnicotinamide, with insulin resistance’, Diabetologia, 58(4), pp. 799–808.

36. Kim, T. N. and Choi, K. M. (2013) ‘Sarcopenia: definition, epidemiology, and pathophysiology’, J Bone Metab, 20(1), pp. 1–10. doi: 10.11005/jbm.2013.20.1.1.

37. Larsen, S. et al. (2012) ‘Biomarkers of mitochondrial content in skeletal muscle of healthy young human subjects’, The Journal of Physiology, 590(14), pp. 3349–3360.

38. Lauretani, F. et al. (2003) ‘Age-associated changes in skeletal muscles and their effect on mobility: an operational diagnosis of sarcopenia’, Journal of Applied Physiology, 95(5), pp. 1851–1860.

39. Lin, A. L. et al. (2015) ‘Caloric restriction increases ketone bodies metabolism and preserves blood flow in aging brain’, Neurobiology of Aging. Elsevier Inc, 36(7), pp. 2296–2303.

40. Liu, L. et al. (2018) ‘Quantitative Analysis of NAD Synthesis-Breakdown Fluxes’, Cell Metabolism, 27(5), p. 1067–1080.e5.

41. Liu, M. et al. (2015) ‘Serum N 1 -Methylnicotinamide Is Associated With Obesity and Diabetes in Chinese’, The Journal of Clinical Endocrinology & Metabolism, 100(8), pp. 3112– 3117.

42. Lu, M. et al. (2014) ‘Intracellular redox state revealed by in vivo 31 P MRS measurement of NAD + and NADH contents in brains’, Magnetic Resonance in Medicine, 71(6), pp. 1959– 1972.

43. Martens, C. R. et al. (2018) ‘Chronic nicotinamide riboside supplementation is well-tolerated and elevates NAD+ in healthy middle-aged and older adults’, Nature Communications, 9(1), p. 1286.

44. Massudi, H. et al. (2012) ‘Age-associated changes in oxidative stress and NAD+ metabolism in human tissue’, PLoS ONE, 7(7), pp. 1–9.

45. Mayer, U. (2003) ‘Integrins: redundant or important players in skeletal muscle?’, The Journal of biological chemistry, 278(17), pp. 14587–90.

46. McNally, E. (2012) ‘Novel Targets and Approaches to Treating Skeletal Muscle Disease’, in Muscle, pp. 1095–1103.

47. Mills, K. F. et al. (2016) ‘Long-Term Administration of Nicotinamide Mononucleotide Mitigates Age-Associated Physiological Decline in Mice’, Cell Metabolism. Elsevier Inc., 24(6), pp. 795–806.

48. Mootha, V. K. et al. (2003) ‘PGC-1α-responsive genes involved in oxidative phosphorylation are coordinately downregulated in human diabetes’, Nature Genetics, 34(3), pp. 267–273.

49. Mouchiroud, L. et al. (2013) ‘The NAD(+)/Sirtuin Pathway Modulates Longevity through Activation of Mitochondrial UPR and FOXO Signaling’, Cell, 154(2), pp. 430–441.

50. Pesta, D. and Gnaiger, E. (2012) ‘High-Resolution Respirometry: OXPHOS Protocols for Human Cells and Permeabilized Fibers from Small Biopsies of Human Muscle’, in Methods in Molecular Biology, pp. 25–58.

51. Phielix, E. et al. (2008) ‘Lower intrinsic ADP stimulated mitochondrial respirationin underlies in vivo mitochondrial dysfunction in muscle of male type 2 diabetic patients’, Diabetes, 57(11), pp. 2943–2949.

52. Polzonetti, V. et al. (2012) ‘Population variability in CD38 activity: Correlation with age and significant effect of TNF-α-308G > A and CD38 184C > G SNPs’, Molecular Genetics and Metabolism, 105, pp. 502–507.

53. Porter Starr, K. N. and Bales, C.W. (2015) ‘Excessive Body Weight in Older Adults’, Clinics in Geriatric Medicine, 31(3), pp. 311–326.

54. Ratajczak, J. et al. (2016) ‘NRK1 controls nicotinamide mononucleotide and nicotinamide riboside metabolism in mammalian cells’, Nature Communications, 7, p. 13103.

55. Ryu, D. et al. (2016) ‘NAD+ repletion improves muscle function in muscular dystrophy and counters global PARylation’, Science Translational Medicine, 8(361), pp. 1–15.

56. Singh, T. and Newman, A. B. (2011) ‘Inflammatory markers in population studies of aging’, Ageing Research Reviews, 10(3), pp. 319–329.

57. Sousa, A. S. et al. (2016) ‘Financial impact of sarcopenia on hospitalization costs.’, European Journal of Clinical Nutrition, 70(9), pp. 1046–1051.

58. Subramanian, A. et al. (2005) ‘Gene set enrichment analysis: A knowledge-based approach for interpreting genome-wide expression profiles’, Proceedings of the National Academy of Sciences, 102(43), pp. 15545–15550.

59. Trammell, S. A. et al. (2016) ‘Nicotinamide riboside is uniquely and orally bioavailable in mice and humans’, Nature Communications, 7, p. 12948.

60. Trammell, S. A. and Brenner, C. (2013) ‘TARGETED, LCMS-BASED METABOLOMICS FOR QUANTITATIVE MEASUREMENT OF NAD+ METABOLITES’, Computational and Structural Biotechnology Journal. Elsevier, 4(5).

61. Trammell, S.A.J. et al. (2016) ‘Nicotinamide Riboside Opposes Type 2 Diabetes and Neuropathy in Mice’, Scientific Reports. Nature Publishing Group, 6(May), p. 26933.

62. Vannini, N. et al. (2019) ‘The NAD-Booster Nicotinamide Riboside Potently Stimulates Hematopoiesis through Increased Mitochondrial Clearance’, Cell Stem Cell, 24(3), p. 405– 418.e7.

63. Vaur, P. et al. (2017) ‘Nicotinamide riboside, a form of Vitamin B3, protects against excitotoxicity-induced axonal degeneration’, FASEB Journal, 31(12), pp. 5440–5452.

64. van de Weijer, T. et al. (2015) ‘Evidence for a direct effect of the NAD+ precursor acipimox on muscle mitochondrial function in humans’, Diabetes, 64(4), pp. 1193–1201.

65. Winter, J. E. et al. (2014) ‘BMI and all-cause mortality in older adults: a meta-analysis’, The American Journal of Clinical Nutrition, 99(4), pp. 875–890.

66. Wythe, S. et al. (2015) ‘Getting the most from venous occlusion plethysmography: Proposed methods for the analysis of data with a rest/exercise protocol’, Extreme Physiology and Medicine, 4(1), pp. 4–8.

67. Yoshino, J., Baur, J. A. and Imai, S.ichiro (2017) ‘NAD+Intermediates: The Biology and Therapeutic Potential of NMN and NR’, Cell Metabolism, 27(3), pp. 513–528.

68. Zhang, H. et al. (2016) ‘NAD+ repletion improves mitochondrial and stem cell function and enhances life span in mice’, Science, 352(6292), pp. 1436–1443.

69. Zhou, C.-C. et al. (2016) ‘Hepatic NAD + deficiency as a therapeutic target for non-alcoholic fatty liver disease in ageing’, British Journal of Pharmacology, 173(15), pp. 2352–2368.

70. Zhu, X.-H. et al. (2015) ‘In vivo NAD assay reveals the intracellular NAD contents and redox state in healthy human brain and their age dependences.’, Proceedings of the National Academy of Sciences of the United States of America, 112(9), pp. 2876–81.

